# Senescence-induced reparative fibroblasts enable scarless wound healing in aged murine skin

**DOI:** 10.1101/2025.04.17.648896

**Authors:** Dongyang Wang, Zhanqi Wang, Yang Yang, Fuwei Bai, Haiyun Li, Junning He, Feng Zhou, Tao Chen, Tongfeng Fang, Zhongyu Wu, Junhao Zhong, Lin Xiang, Yi Man, Yingying Wu

## Abstract

It has been reported that elderly individuals exhibit reduced scarring during the wound healing process compared to younger adults. However, the underlying mechanisms responsible for this phenomenon remain poorly understood. Here, we revealed that aged mice exhibited more pronounced regenerative outcomes compared to young mice, characterized by increased hair follicle numbers and collagen fiber features closer to normal skin. Single-cell sequencing identified a reparative fibroblast subpopulation (Prss35^+^Fib) enriched in the aged group, which promotes regeneration through communication with epithelial cells, macrophages, and T cell subpopulations via PTN and EREG signaling. Spatial transcriptomics validated this communication pattern by elucidating cell proximity and locating the regenerative niche in the upper dermis. Finally, EREG treatment significantly enhanced regenerative outcomes in young mice, while the small wound model of aged mice, lacking the reparative fibroblast and EREG signaling, failed to achieve regeneration. Collectively, our findings advance the understanding of regenerative plasticity in aging and provide new insights for designing scarless therapeutic strategies.

## Introduction

Two distinct repair patterns emerge following severe cutaneous injury: regenerative or fibrotic healing^1^. Regenerative healing reconstructs native tissue architecture and function, entailing de novo hair follicle (HF) and physiological extracellular matrix (ECM)^2,3^. Yet fibrotic healing replaces functional dermis with disorganized ECM deposition, culminating in scar or keloid formation, which profoundly compromises patients’ physical function and psychosocial well-being^4–6^. Although numerous studies have drafted the mechanism of regenerative and fibrotic healing, the challenges to clinical translation remain unsolved^2,7–9^. However, natural regenerative programs may hold therapeutic promise, as minimal scarring has been observed in elderly individuals compared to adults^10^. Given age-related dysfunctions (impaired angiogenesis and persistent inflammation), the attenuated velocity of wound healing with advancing age is well-documented, while the mechanism of minimal scarring is rarely reported^11,12^. Recent work demonstrated that aged rodents exhibit reduced scarring in ears’ wound models, with deficiency of stromal cell-derived factor 1 enhancing full-thickness tissue regeneration^13^. As the performance of secreted factors found is only part of scar formation, the regenerative potential of aged individuals needs further investigation.

Current research reveals the physiological role of cellular senescence in regeneration across species^14^. In fin amputation models of zebrafish, injury-induced senescent cells promote tissue regeneration, though the mechanism remains unclear^15^. Further studies in cnidarian and newts showed that senescent cells reprogram neighboring cells into the pro-regenerative state to facilitate tissue regeneration, probably through senescence-associated secretory phenotype (SASP)^16,17^. While this mechanism remains understudied in mammalian skin, we propose that aging-related accumulation of senescent cells and SASP factors may enable injury-site cells to develop analogous reprogramming capacities, thereby orchestrating reparative functions. As key signaling senders that organize anatomical niches in skin repair, fibroblasts critically mediate the balance between regenerative and fibrotic healing processes^18^. During the remodeling of scarring, the activated fibroblasts differentiate into heterogeneous lineages, including fibrogenic and regenerative state^19,20^. Previous work discovered that fibrogenic lineages can transform into other mesenchymal cells during wound healing, suggesting the reprogramming potential of fibroblast in mammalian skin^21,22^. While the heterogeneity of fibroblast has been elucidated, critical gaps persist in understanding whether reprogramming of fibroblast lineages emerges and through which molecular mediators govern the scarless healing of aged skin ^11,23^.

In this study, we compared wound healing patterns between aged and young mice following full-thickness skin injury. Through integrated single-cell RNA sequencing (scRNA-seq) and spatial transcriptomics (ST), we identified a unique reparative fibroblast subpopulation in aged mice that reprograms from fibrogenic to regenerative states, a process linked to cellular senescence. Histological analyses further revealed these fibroblasts coordinate regenerative niches through specific SASP factors. By validating findings with published datasets and in vivo functional assays, we confirmed the conserved role of this regenerative mechanism.

## Result

### 3.1 Wound healing in aged skin was approaching a regenerative pattern

To investigate age-dependent healing patterns, we established a splinted excisional wound model on the dorsal skin of young (6-week-old) and aged (18-month-old) mice (Fig. 1A). This model was selected based on its validated capacity to inhibit wound contraction, thereby recapitulating human re-epithelialization dynamics^24^. Then three phases of the healing process were stratified and performed subsequent analyses^25^: coagulation (7 days post-wounding, dpw), inflammation (14 dpw), and scar formation (21&28 dpw)(Supplementary Fig. 1A). Consistent with aging-related healing impairments, aged mice exhibited significantly delayed wound closure at 14 dpw (Supplementary Fig. 1B, p < 0.01), a phenomenon not further investigated due to our prior research^26^.

**Fig 1:**
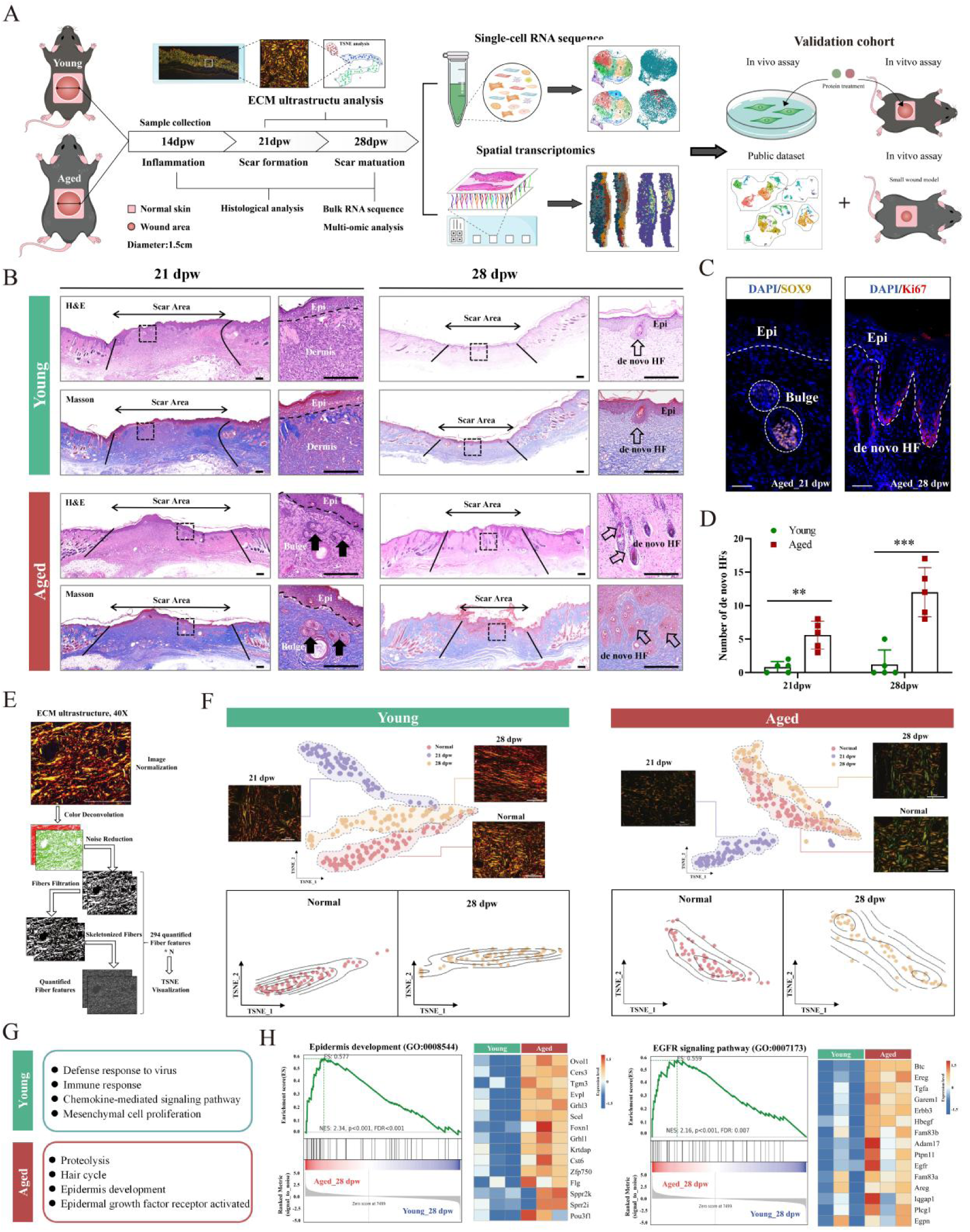
Wound healing in aged skin was approaching a regenerative pattern. (A) Schematic diagram of study design, multi-omics analysis, and validation cohort. (B) Hematoxylin-eosin (H&E) and Masson staining for scar tissues, displaying morphological differences between the young and aged groups. Scale bar, 200 um. (C) Immunofluorescence staining for Sox9 (marker of the bulge) and Ki67 (marker of de novo HF) in the scar tissues of the aged group. Scale bar, 50um. (D) Quantification analysis for numbers of de novo HFs (n=5 per group). Significances were analyzed using bidirectional t-tests. (E-F) Diagram of ECM ultrastructure analysis(E). TSNE plots of unsupervised cluster analysis of ECM characteristics in each group with representative Sirius red staining images. Scale bar, 50um. (F-H) Bulk RNA sequencing for scar tissues at 28 dpw(n=3 per group). Gene ontology enrichment analysis showing the top biological process terms based on differentially expressed genes (p<0.05, |log2FC|>1)(G). Gene set enrichment analysis (GSEA) showing regeneration-associated terms (left) and heatmap of corresponding genes (right)(H). Abbreviation: dpw, day post wound. HF, hair follicle. Epi, epidermis. ECM, extracellular matrix. TSNE, T-Distributed Stochastic Neighbor Embedding. EGFR, epidermal growth factor receptor. Statistical thresholds followed p = 0.05 convention (*p < 0.05; **p < 0.01; ***p < 0.001; ****p < 0.0001), with nonsignificant (ns) determinations requiring p > 0.05.

However, we unexpectedly found that aged mice demonstrated reduced scar formation with regenerative features at 28 dpw, a previously understudied late-phase healing phenotype (Fig.1B). Using the non-regenerative panniculus carnosus as an anatomical boundary, we distinguished scar tissue from uninjured dermis^24^. Histological analysis revealed prototypical scarring in young mice, whereas aged scars exhibited regeneration of the pilosebaceous unit. Bulge structures emerged at 21 dpw, progressing to mature de novo hair follicles (HFs) by 28 dpw. Immunofluorescence (IF) for SOX9 (hair follicle stem cell marker) and Ki-67 (proliferation marker)^2^ confirmed active epithelial proliferation in aged de novo HFs (Fig. 1C-D). Quantification demonstrated the number of de novo HFs in 21dpw (mean=5.6, p<0.01) and 28dpw (mean=12, p<0.001) in the aged group was significantly higher than that in the young group (21dpw, mean=0.8; 28dpw, mean=1.2).

Given the association between scar maturation and collagen reorganization^25^, the architecture of the extracellular matrix (ECM) in the scar area was further analyzed. Both groups exhibited disorganized collagen at 21 dpw (characteristic of immature scars). Notably, aged mice maintained physiologic collagen ratios (p = 0.5745 vs normal skin) at 28 dpw, sharply contrasting with the dense collagen bundles in young mice (Supplementary Fig. 1C). Then we employed a newly validated ECM ultrastructure algorithm to quantify analysis parameters of the ECM^19,20^, including fibril length and angular distribution, etc (Supplementary Table 1). T-distributed stochastic neighbor embedding (TSNE) clustering demonstrated progressive normalization of aged ECM architecture, with 28 dpw samples clustering closer to uninjured skin than young counterparts (Fig. 1E-F). Sirius red stain revealed type I collagen predominance in the young group, versus physiological type I/III collagen in the aged (Supplementary Fig. 1D).

To further verify the regenerative healing ability of the aged group, Bulk RNA sequencing was performed on the scar tissues at 28 dpw, Gene ontology (GO) and Gene set enrichment analysis (GSEA) were performed for differentially expressed genes (p<0.05,|log2FC|>1). The young group upregulated mesenchymal cell proliferation and immune response (established pro-fibrotic drivers)^2^; while the aged group activated epithelial development, hair follicle cycle, and epidermis growth factor receptor (EGFR) signaling (closely related to re-epithelialization and hair growth) (Fig. 1G-H). For instance, amphiregulin (Areg), as an EGFR ligand, has been proven to be a key effector in promoting HF regeneration in previous studies^27,28^.

Collectively, regeneration of de novo HFs, normalization of the ECM, and pro-regenerative transcriptional signatures support scarless healing in the aged mice, though the specific mechanisms require further elucidation.

### 3.2 Single-cell profiling reveals heterogeneity underlying age-specific clusters

To delineate cellular determinants of the aged regenerative phenotype, we performed scRNA-seq on scar tissues at 28 dpw, guided by our prior observation of aged scars resembling uninjured skin architecture. Following quality control, we captured 12,510 cells from the young group and 8,638 from the aged group. Uniform manifold approximation and projection (UMAP) clustering coupled with canonical gene marker analysis identified major cell types (Fig. 2A, Supplementary Fig. 2A-B), revealing compositional differences: young scars contained higher proportions of fibroblasts (76% in young vs 58% in aged) and epithelial cells (6% vs 4%), whereas aged scars were enriched with macrophages (2% vs 8%) and T/NK cells (4% vs 13%).

**Fig 2:**
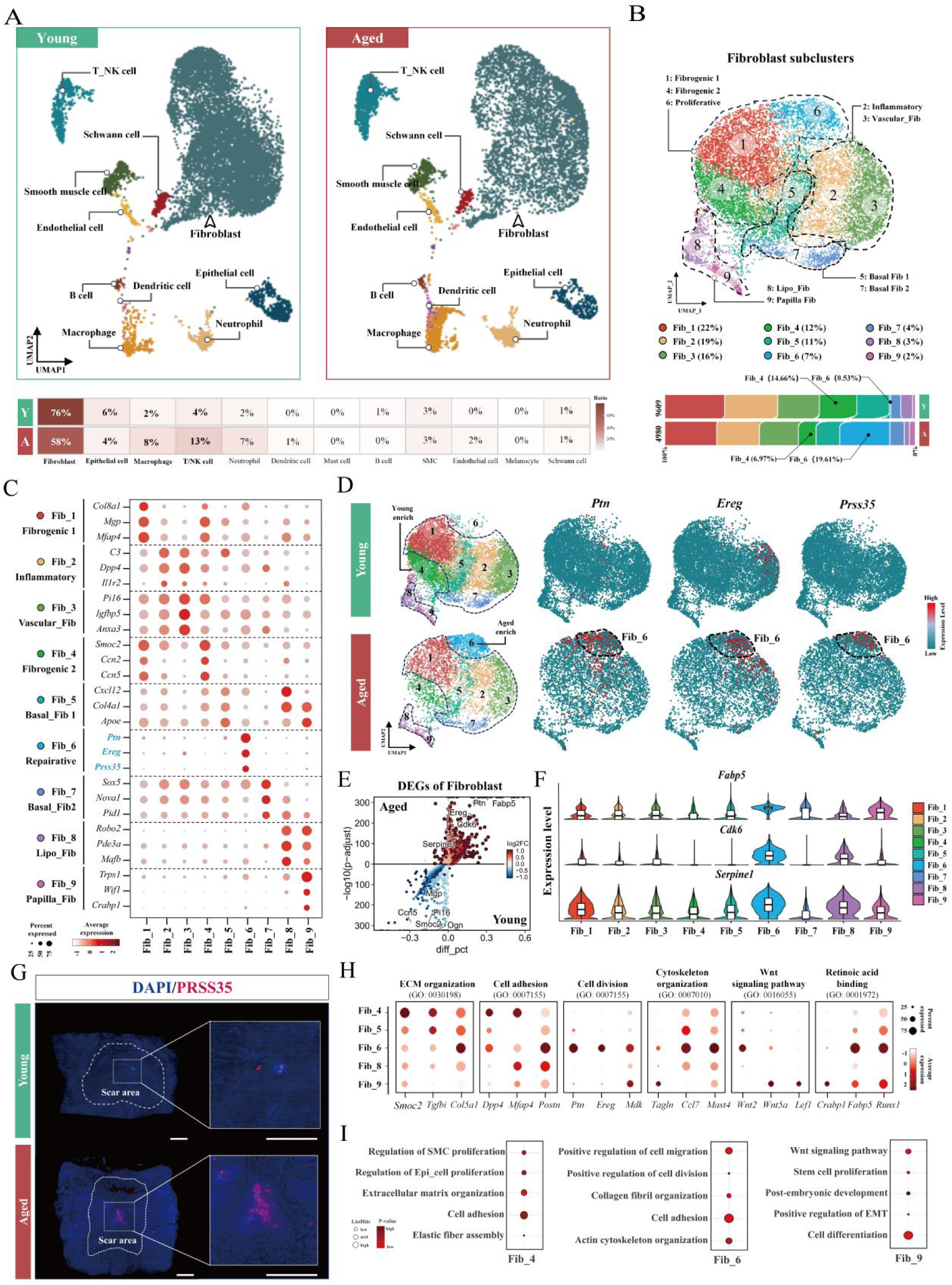
Single-cell profiling reveals heterogeneity underlying age-specific clusters. (A) UMAP plots showing the proportion and distribution of major cell types from scar tissues. (B) UMAP plot showing nine subclusters of fibroblast with their proportions. Distinct Color indicates the functional identity of subclusters. Y (Young) and A (Aged). (C-D) Dotplot showing marker genes of the fibroblast subclusters(C). Feature plots showing age-specific subcluster and marker genes(D). (E) Tissue clearing staining for Prss35 (red, marker gene of Fib_6) in scar tissues at 28 dpw. The dotted line distinguished the scar area from uninjured skin. Scale bar, 500um. (F) Volcano plot showing DEGs of fibroblast within groups based on Wilcoxon rank-sum test (left). Vlnplot showing expression level of aged-specific genes within fibroblasts subclusters(right). (G-H) Dot plot showing expression profile of fibrotic and regenerative marker genes (G) and GO enrichment terms (H) within characteristic fibroblast subclusters. Abbreviation: DEGs, differentially expressed genes. GO, gene ontology. SMC, smooth muscle cell. Epi_cell, epithelial cell. EMT, epithelial-mesenchymal transition.

Given fibroblasts’ established involvement in collagen remodeling, immune modulation, and epidermal reprogramming, we first conducted the sub-clustering analysis to deconstruct their functional heterogeneity^8^. Nine transcriptionally distinct fibroblast subclusters were identified (Fig. 2B-D), including basal Fib, fibrogenic Fib, inflammatory Fib, and papillary Fib. The comparative analysis uncovered age-specific distribution patterns: Fib_6, nearly exclusive to aged scars (0.53% in young vs 19.61% in aged), contrasted with Fib_4 predominance in the young group (14.66% vs 6.97%). Notably, Fib_6 exhibited marked enrichment of regenerative-associated markers, including *Ptn*, *Ereg*, and *Prss35*, suggesting a pro-reparative phenotype^29,30^. Spatial mapping via tissue clearing stain confirmed *Prss35* (the marker gene of Fib_6) predominantly localized to scar centers in the aged group (Fig. 2E), mirroring the spatial distribution of de novo HFs (Fig. 1B), consistent with its reparative definition.

Further analysis of fibroblasts revealed distinct age-related gene expression profiles^31^. Fib_4 (predominant in the young group) upregulated pro-fibrotic genes (*Smoc2*)^31^, while Fib_6 (aged-specific cluster) displayed cell cycle progression (*Cdk6*, *Serpine1*)^32^, supporting their characteristic designation subsequent analysis (Fig. 2F). Comparative analysis revealed Fib_6’s differential co-expression of fibrotic (*Postn*) and regenerative markers (*Fabp5*, *Runx1*), with unique *Wnt2* expression, a key developmental signaling molecule in human skin (Fig. 2G)^8,33^. GO enrichment analysis demonstrated subset functional specialization: Fib_4 mediated ECM remodeling/elastic fiber assembly (collaborating with Fib_1 [*Mgp^+^/Mfap4^+^*] in the fibrotic healing)^31,34^; Fib_6 activated cellular proliferation/migration capacity; Fib_9 regulated Wnt signaling pathway in the dermal papilla formation^35^ (Fig. 2H, Supplementary Fig. 2C).

Epithelial sub-clustering identified hair follicle progenitors *(Lef1^+^*), basal (*Krt14^+^*), and spinous (*Krt1^+^*) clusters, with aged mice showing increased progenitor populations consistent with enhanced HF regeneration (Supplementary Fig. 2D). Immune subclusters analysis revealed aged-specific expansion of anti-inflammatory macrophages(*Arg1^+^*), Th17 and regulatory T cells (Treg), consistent with our previous finding in immune micro-environment during wound regeneration^24^ (Supplementary Fig. S2E-F). These observations uncovered age-specific fibroblast and immunomodulatory shifts, while the molecular mechanisms orchestrating their reparative potential and spatial interplay remain undefined.

### 3.3 Spatial distribution and differentiated trajectory of reparative fibroblast

Given the central localization of neo-HFs in the aged group and spatial congruence of Fib_6 (*Prss35^+^*), we applied ST to explore further the spatial heterogeneity of scar tissues at 28dpw (Fig. 1A). Serial sections with H&E staining and imaging identified sections containing maximal scar regions for ST. Post-quality control (Supplementary Fig. 3A), molecular feature-based clustering partitioned spatial spots into distinct clusters: cluster 3 (epidermis), cluster 4 (dermis), clusters 1-2 (hypodermis) (Fig. 3A, Supplementary Fig. 3B-C). To map the single-cell data to the spatial level, we employed the RCTD algorithm^36^ to anchor the spatial distribution of cell subpopulations (Fig. 3B). AddModuleScore-based fibroblast mapping demonstrated spatial stratification: pro-fibrotic Fib_1/Fib_4 localized to hypodermal layers, versus pro-regenerative Fib_6/Fib_9 restricted to the papillary dermis. Quantitative analysis demonstrated young mice exhibited higher Fib_1/Fib_4 proportions (p < 0.0001), while Fib_6 (p < 0.01) and Fib_9 (p < 0.05) showed aged-specific accumulation in central scar regions (Fig. 3C-D, Supplementary Fig. S3D). This echoes prior research that wound-induced regenerative fibroblasts (*Crabp1^+^/Fabp5^+^/Runx1^+^*) exclusively occupy the central upper dermis in HF regeneration models, implying spatial positioning dictates regenerative capacity^8^.

**Fig 3:**
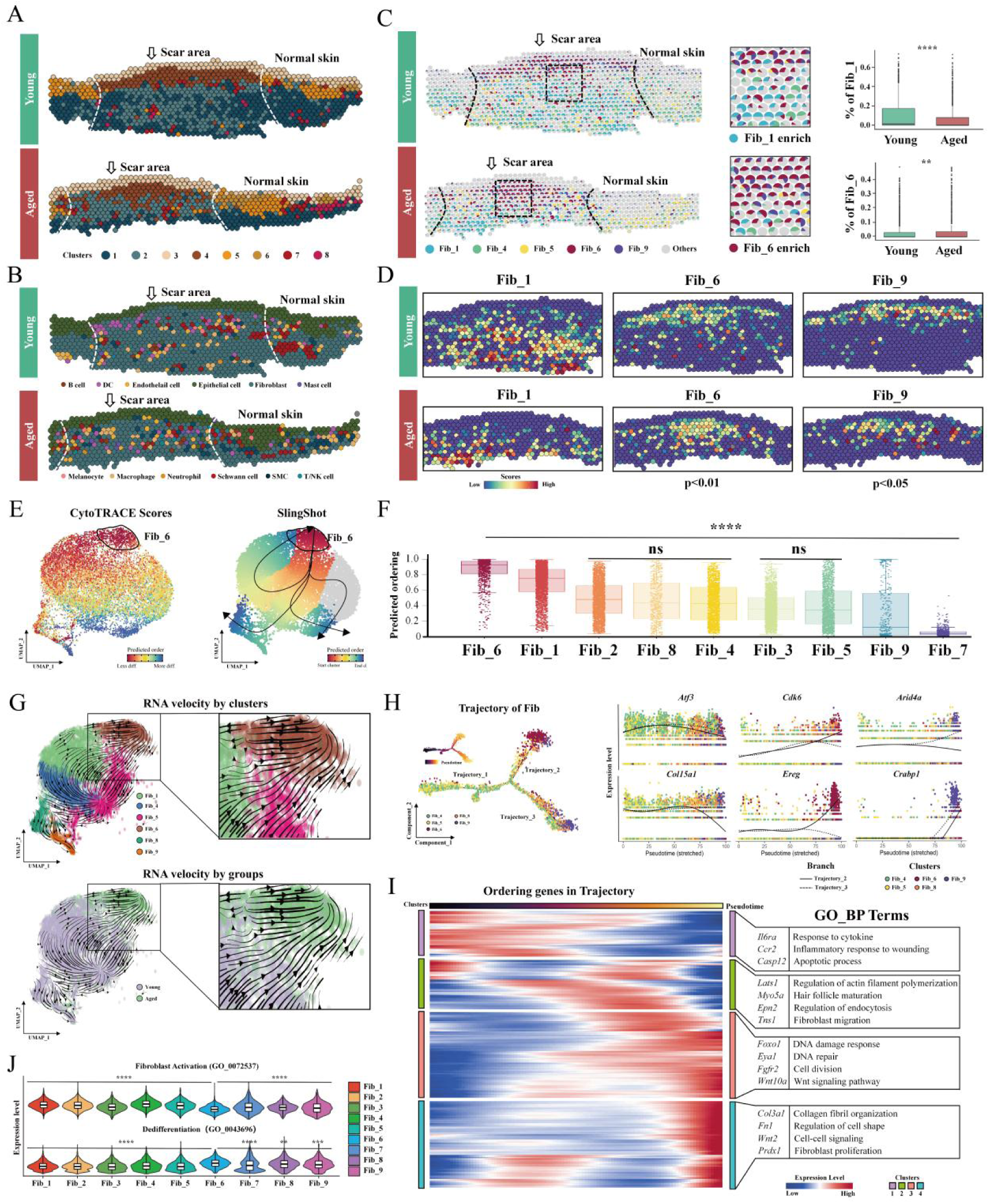
Spatial distribution and differentiated trajectory of reparative fibroblast. (A-B) Spatial map showing unsupervised cluster (A) and major cell types based on RCTD algorithm (B) in spatial transcriptomics. The dotted line distinguished the scar area from normal skin. (C) Spatial pie plot showing the spatial stratification of fibroblast subclusters (left). Box plot showing the difference in the proportion of subclusters using the Wilcoxon test (right). (D) Spatial feature plots showing the distribution characteristics and proportion difference analyzed by the Wilcoxon test of fibroblast subclusters. (E) UMAP plots showing CytoTRACE scores (left) and differentiation trajectory inferred by SlingShot (right) of fibroblast subclusters. Arrows indicate the direction of differentiation. (F) Box plot showing predicted ordering and significance difference of fibroblasts. The multigroup comparisons against the control group (Fib_6) employed one-way ANOVA with Dunnett’s post hoc analysis. (G) UMAP plots showing the projection of RNA velocity, with arrows indicating the direction of differentiation. (H) Monocle2 pseudotime analysis of the trajectory map and branch lines labeled the changing trend of key gene expression. (I) Heatmap showing ordering genes in pseudotime trajectory with GO enrichment terms. (J) Vlnplot showed the gene set scores compared with Fib_6 in the multi-group comparison. For statistical analysis methods, see (F). Abbreviation: GO_BP terms, gene ontology_biological process terms. Statistical thresholds followed p = 0.05 convention (*p < 0.05; **p < 0.01; ***p < 0.001; ****p < 0.0001), with nonsignificant (ns) determinations requiring p > 0.05.

We subsequently applied multiple pseudotemporal analysis algorithms to investigate Fib_6’s position in differentiation trajectories. Using CytoTRACE to quantify gene expression per cell^37^, we assessed fibroblast subcluster differentiation states and developmental potential (Fig. 3E-F). The analysis predicted Fib_6 as having the highest developmental potential (p<0.0001), supporting its designation as the differentiation origin. Surprisingly, Slingshot trajectory construction suggested Fib_6 served as a differential origin and endpoint, a paradox potentially arising from its transitional state. To resolve this paradox, we performed RNA velocity analysis^38^. By calculating spliced/unspliced RNA ratios, this method reconstructs transcriptional dynamics during differentiation (Supplementary Fig. 4A-B). RNA velocity vector mapping on UMAP plots identified Fib_5 as the primary differentiation origin, with Fib_6 receiving inputs from Fib_5, Fib_4, and Fib_1 (Fig. 3G). Given Fib_4 and Fib_1’s fibrogenic roles, we propose Fib_6 as in a transitional state between fibrogenic and regenerative, which aligns with its hybrid expression profile and explains its ambiguous positioning in prior analyses. Prior studies highlight fibroblast plasticity and heterogeneity^18,39^, suggesting myofibroblasts may transition between activation states. However, whether reparative Fib directly originates from pro-fibrotic subtypes remains unclear, necessitating lineage tracing for further validation.

Monocle2 analysis identified the driver genes of the above differentiation trajectory (Fig. 3H). Pseudotemporal ordering revealed progressive downregulation of fibrotic genes (*Atf3*, *Col15a1*) in Trajectories 2-3, while Trajectory_2 upregulated regenerative markers (*Cdk6, Ereg, Crabp1*), and Trajectory_3 activated proliferation-related genes (*Arid4a*)^40,41^. RNA velocity analysis corroborated these findings, showing Fib-1/Fib-4 driven by the pro-fibrotic gene (*Fam114a1*) and Fib-9 by the Wnt signaling-associated gene (*Tmeme59*) (Supplementary Fig. 4E)^42,43^. Pseudotime-dependent GO enrichment analysis demonstrated sequential activation: cellular migration (*Tns1*), Wnt signaling (*Wnt10a*), and morphogenesis regulation *(Fn1, Prdx1*) (Fig. 3I). Gene set scoring confirmed Fib_6’s quiescent state with elevated dedifferentiation markers, collectively supporting its transition from fibrotic to regenerative states (Fig. 3J).

Notably, DNA damage response (*Foxo1*) and repair pathways emerged as key pseudotemporal features (Fig. 3I)^44^. RNA velocity further identified Fib_6-specific drivers *Rsf1* (DNA repair) and *Ereg* (SASP factor), suggesting DNA damage-induced senescence may prime Fib_6 for reprogramming with the acquisition of regenerative potential (Supplementary Fig. 4E-F). Although senescence is classically considered regeneration-inhibitory, emerging evidence suggests senescent cells exhibit enhanced plasticity for reprogramming and tissue regeneration ^14–17,45,46^. Comparative analysis exhibited feature genes of cellular senescence in reparative fibroblast-Fib_6 (*Areg, Bmp6, Ereg, Fgf7, Mif, Serpine1, Spp1*) (Supplementary Fig. 4G).

Our integrated multi-omics analyses demonstrate that the aged scar microenvironment orchestrates two key processes: (1) the reduction of fibrogenic fibroblasts and (2) senescence-mediated reprogramming of reparative fibroblasts, which are spatially confined to regenerative regions.

### 3.4 Difference of cellular communication patterns of reparative fibroblast

To explore the molecular driver’s reparative functions of age-specific clusters, we conducted cell-cell crosstalk analysis subsequently. Reparative fibroblast (Fib_6) revealed enhanced signaling activity, with both incoming and outgoing interaction strength surpassing other cell subsets (Fig. 4A, Supplementary Fig. 5A). Notably, Fib_6 and papillary Fib_9 engaged unique regenerative signaling axes, including *PTN, EGF, MDK*, and non-canonical Wnt (ncWNT) pathways^2^, mirroring global communication shifts where aged scars favored regenerative signals (PTN/EGF) versus young fibrotic signatures (TGF-β) (Fig. 4B, Supplementary Fig. 5B-D). Ligand-receptor visualization identified Fib_6 as the dominant sender of PTN and EGF signals, though MDK and TGF-β signaling lacked this specificity (Supplementary Fig. 5E). These findings suggest Fib_6’s reparative capacity stems from the selective expression of pro-regenerative ligands, particularly PTN (a conserved niche organizer in tissue regeneration and folliculogenesis^47^) and EREG (recently implicated in HF restoration^48^). Focusing on fibroblast-epithelial crosstalk, Fib_6 demonstrated robust PTN/EREG signaling to epithelial cells and fibroblast subsets (Fib_4/Fib_9) (Fig. 4C), contrasting with Fib_4’s TGF-β-dominated profile. Intriguingly, Fib_6 also secreted POSTN, classically a myofibroblast marker, which paradoxically marked regenerative fibroblasts in reindeer and human wound models^2^. Though murine *POSTN+* fibroblast biology remains underexplored, we postulate its dual role may facilitate Fib_6’s functional plasticity^49^.

**Fig 4:**
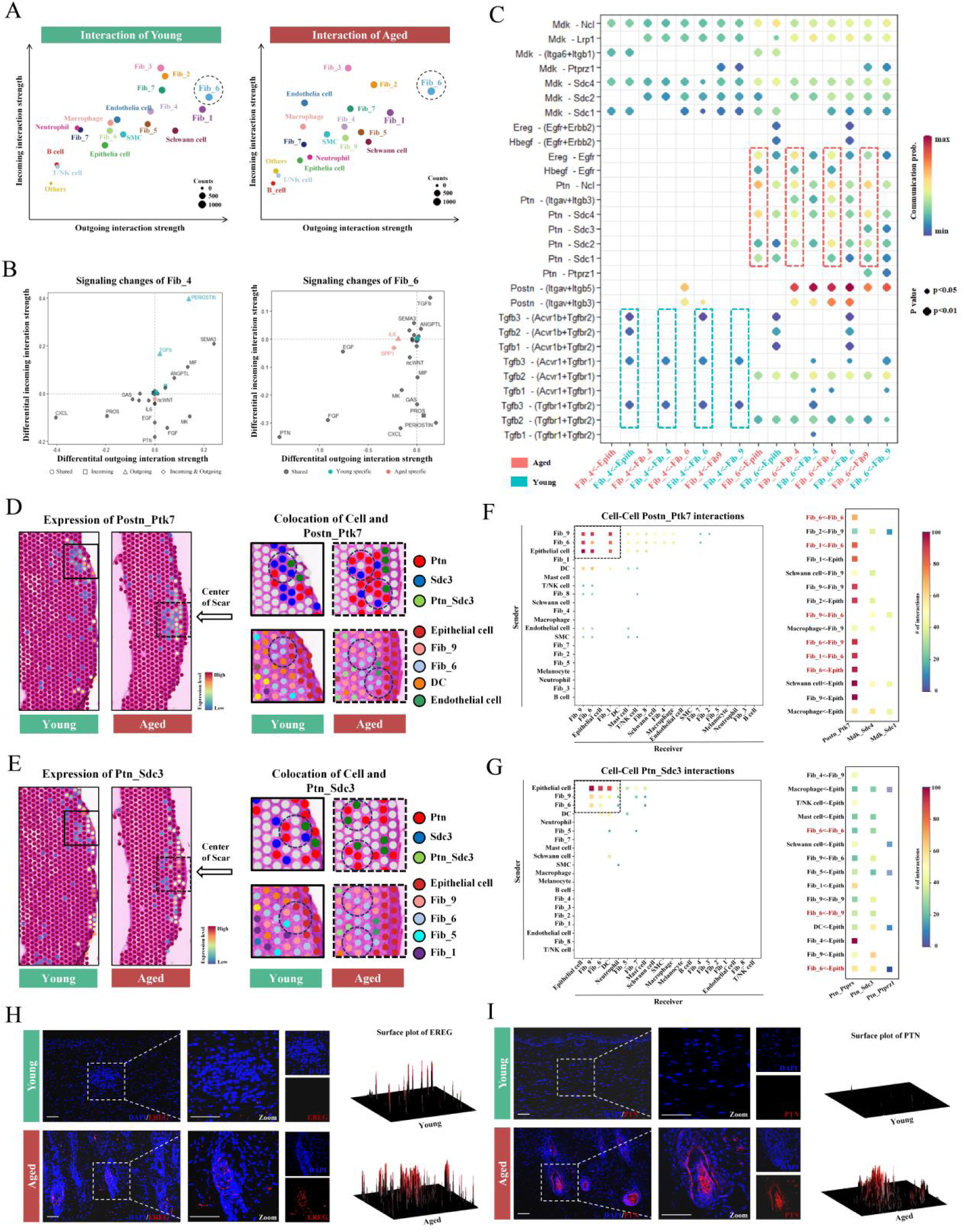
Difference of cellular communication patterns of reparative fibroblast. (A) Integrated incoming and outgoing cell-cell interaction strength using cellchat. The counts represented the total number of interactions per subcluster across all signaling pathways. (B) The Signaling changes of characteristic fibroblasts between the young and aged groups, with shapes and colors identifying the type of signaling changes. The upper right and lower left quadrants represented up-regulated signaling pathways in the young group compared with the aged, respectively. (C) Bubble plot of age-specific ligand and receptor pairs between fibroblasts and epithelial cells. The blue dotted box represented young specific pairs and the red represented aged specific. (D-E) Spatial feature plot showing the expression level of LR pairs using the stlearn algorithm. The color represents whether an LR pair interacts significantly more strongly than a random background at a particular spot. (left). The solid line square represented zoom images of the young group and the dotted line square represented the aged. Dotted circles signed the colocation of cell type and ligand-receptor pairs (right). (F-G) The strength of cell-cell interactions (left) and top cell-cell pairs in specific signaling(right) in ST was assessed by permutation test (n_perms=1000). (H-I) Immunofluorescence staining for EREG (H) and PTN (I) in the scar tissues between young and aged groups. Scale bar, 50um. Surface plot showing relative fluorescence intensity of EREG and PTN in zoom images. Abbreviation: LR, ligand and receptor.

Focusing on regeneration-relevant epithelial subsets (Epith_H: hair follicle-associated; Epith_B: basal Epithelial cell), we identified Fib_6 as the dominant sender of pro-regenerative signals to epithelial cells, including EREG-EGFR, PTN-SDC4, and WNT5A (Fig. 4C, Supplementary Fig. 6A), mechanistically linking to epithelial proliferation and follicular stem cell activation. Reciprocally, epithelial cells promoted fibroblast expansion through FGF signaling. Macrophage subcluster analysis revealed aged-enriched Arg1^+^ macrophages and young-predominant C1qa^+^ macrophages communicating with Fib_6 and Fib_9 via EGF. While the Arg1^+^ subcluster uniquely communicated through SPP1 and MIF (Supplementary Fig. 6B), senescence-associated factors potentially facilitated Fib_6’s fibrotic-to-reparative reprogramming^45^. The interactions of Th17 cells further reinforced regenerative signaling, communicating with fibroblasts via AREG-EGFR (Supplementary Fig. 6C-D), consistent with our prior demonstration in the biomaterial-induced wound regeneration model^28^. Notably, Fib_6-specific IL-17 signaling emerged as a novel axis, potentially synergizing with EGFR to drive re-epithelialization, a mechanism recently implicated in regenerative wound closure^50^.

Finally, we applied cell-cell crosstalk analysis on ST to validate cellular subpopulations and ligand-receptor colocalization at the spatial level^51^. Consistent with our communication analysis, Fib_6 primarily interacted with Fib_9, while Fib_9 and Fib_4 communicated with epithelial cells (Supplementary Fig. 7A). Although most signals represented cell-matrix interactions (e.g., Col1a2-Itgb2), Fib_6-Fib_9 specifically engaged WNT signaling (Wnt5a-Fzd1/2), suggesting their synergistic pro-regenerative role (Supplementary Fig. 7B). Spatial mapping of communication signals revealed age-dependent distributions: POSTN/PTN signals localized to wound peripheries in young mice versus central regenerative regions in aged scars (Fig. 4E-F). Colocalization analysis confirmed POSTN/PTN hotspots overlapped with Fib_6/Fib_9 spatial domains. In addition, epithelial cells served as primary sources of POSTN-PTK7 and PTN-SDC3 signals but lacked corresponding receptor engagement (Fig. 4G-H), potentially due to undetected EGF/FGF expression in spatial transcriptomics(Supplementary Fig. 7C), a technical limitation of single-section analysis compared to whole-tissue scRNA-seq. Immunofluorescence validation of 28 dpw scars demonstrated elevated PTN and EREG expression in aged versus young groups, confirming regenerative signaling activation (Fig 4G).

### 3.5 Spatial proximity of reparative fibroblast and multiple cell populations

Building on the spatial dependency of intercellular communication, we performed niche analysis to delineate the global characteristics of cell proximity^52^. By integrating spots with similar cell composition as a subpopulation and reducing dimension clustering, four cell type niches were defined (Fig. 5A): niche_1 (epidermal) containing epithelial cells and papillary fibroblasts, niche_2 (upper dermal) enriched with quiescent/reparative fibroblasts (Fib 5-9), and niche_3-4 (lower dermal) harboring pro-fibrotic Fib_1/Fib_4^18^. Quantitative niche mapping demonstrated aged-specific expansion of regenerative niches (niche_1-2) with reduction of fibrotic niche_4 (Fig. 5B). Integrated differential gene analysis with functional enrichment delineated niche-specific roles: niche_2 formed a regenerative unit where Fib_6 coordinated with epithelial cells, Fib_7, and Fib_9 to drive ECM remodeling in young mice and Hippo signaling activation in aged counterparts. niche_3-4 constituted a pro-fibrosis unit dominated by fibrogenic and inflammatory fibroblasts, regulating myocyte differentiation/contraction and driving fibroblast migration in young groups, a process mechanistically linked to fascial-layer fibroblast mobilization and scar formation (Fig. 5C). Conversely, aged-specific retinoic acid signaling activation in niche_2 aligned with prior Fib_6 expression characterization.

**Fig 5:**
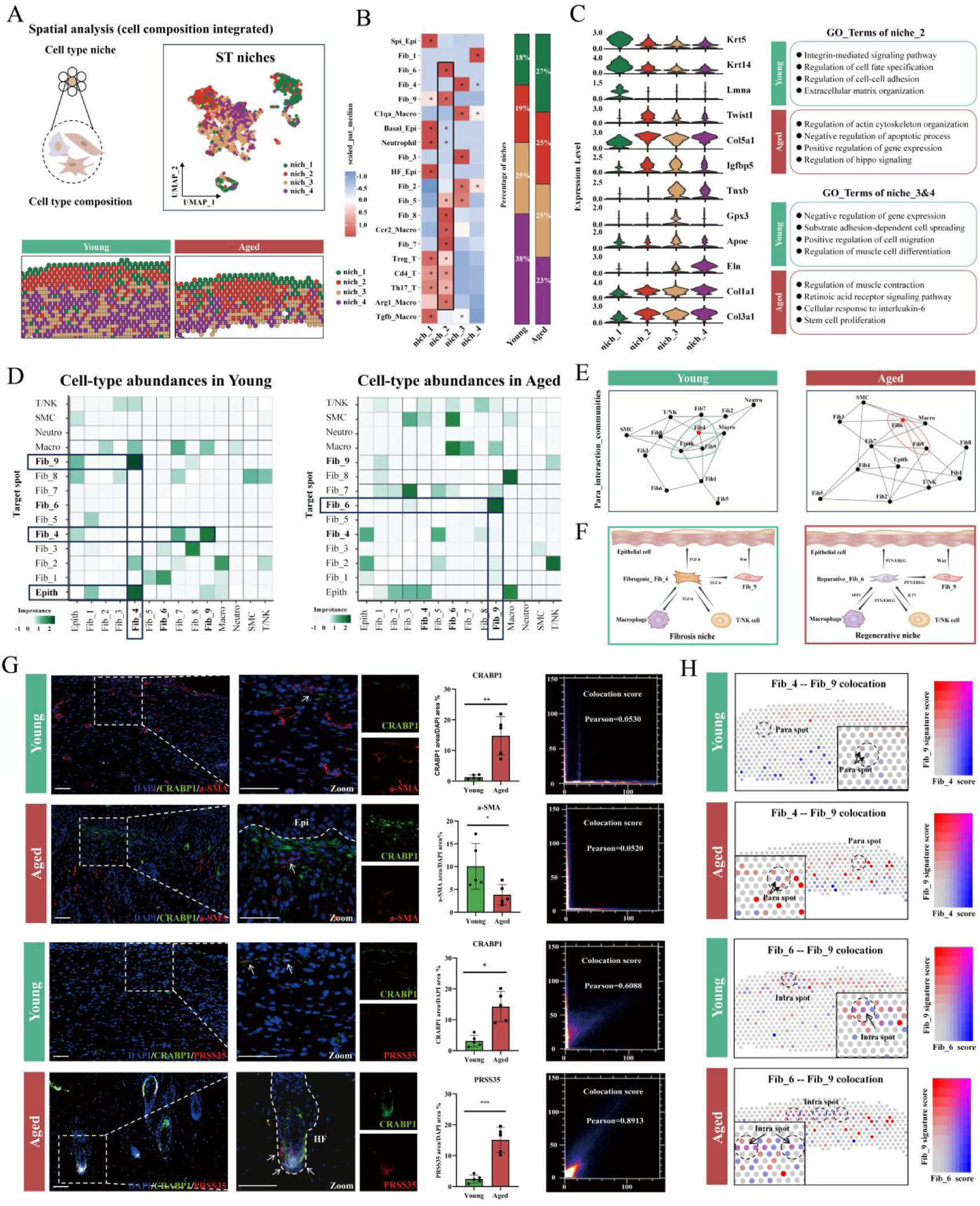
Spatial proximity of reparative fibroblast and multiple cell populations. (A) Cell type niche definition based on integrated cell composition of spots. UMAP plot (upper) and spatial map (down) showing the distribution of four niches. (B) Heatmap plot showing the significance of selected cell types in distinct niches assessed by the Wilcoxon test (left). The percentage of niches between young and aged groups (right). (C) Vlnplot showed top expressed genes in distinct niches (left). The up-regulated GO terms of niches are based on differentially top-expressed genes(right). (D-E) Heatmap plots showing the associations within the para spot in ST using the MistyR algorithm, with the color of importance interpreted as spatial dependency (D). The communities plots showing spatial interactions (gray line) and proximity (polygon) within the para spot (E). (F) Schematic diagram of signaling and spatial proximity of characteristic cells. (G) Immunofluorescence staining for CRABP1 (green, marker of papilla fibroblast), a-SMA (red, marker of myofibroblast), and PRSS35 (red, marker of reparative fibroblast) in the scar tissues between young and aged group(left). Quantification analysis for the ratio of marker protein area/DAPI area (n=5 per group). Significances were utilized using bidirectional t-tests(middle). Colocation plot showing colocalization of CARBP1/a-SMA and CRABP1/PRSS35 assessed by Pearson correlation coefficient using the colocalization finder plugin in ImageJ (right). (H) Spatial feature plot of signature scores showing spatial proximity within fibroblast subclusters. The purple spot represented two subclusters in the intra-spot. Abbreviation: Spi_Epi/Basal_Epi/HF_Epi, spinous/basal/hair follicle progenitors epithelial cells. Macro, macrophage. Cd4_T/Th17_T, Cd4+/Th17 T cells. GO, gene ontology. SMC, smooth muscle cell. Epith, epithelial cell. Neutro, neutrophil. T/NK, T/NK cells. Epi, Epidermis. HF, hair follicle. Statistical thresholds followed p = 0.05 convention (*p < 0.05; **p < 0.01; ***p < 0.001; ****p < 0.0001), with nonsignificant (ns) determinations requiring p > 0.05.

To further explore the spatial relationships of multiple cell populations, we employed MistyR^52^ to analyze nearest-neighbor associations within the para spot and intra spot, with importance interpreted as spatial dependency. The young group exhibited proximity between epithelial cells, Fib_9, and Fib_4, whereas the aged showed Fib_6-Fib_9 adjacency, aligning with niche architecture (Fig. 5D-E). In addition, intra spot analysis revealed Fib_6-Fib_9 co-localization in the young group versus Fib_4-Fib_9 pairing in the aged (Supplementary Fig. 8A-B), suggesting latent regenerative potential in young skin and residual fibrotic activity in aged tissue, a dynamic equilibrium dictating ultimate healing outcomes. Integrated with prior communication results, we mapped group-specific spatial signaling landscapes (Fig. 5F). Immunofluorescence for α-SMA (myofibroblast marker) and CRABP1 (Fib_9 marker) confirmed spatial segregation: α-SMA^+^ fibroblasts dominated in young scars versus CRABP1+ enrichment in aged tissue, with minimal co-localization observed. Using PRSS35 (Fib_6 marker), we detected scarce Crabp1^+^/Prss35^+^ fibroblasts in the young group, contrasting with dual-positive cells within aged de novo HF (Fig. 5G). Notably, young scars at 21 dpw transiently upregulated PRSS35+ fibroblasts (Supplementary Fig. 8C), which potentially mediated response to hyperosmotic stress^53^. Spatial probability scoring of marker genes quantified co-localization patterns: aged scars showed elevated Fib_6-Fib_9 proximity versus comparable Fib_4-Fib_9 para-spot relationships across groups, consistent with IF spatial profiles.

Finally, we delineated spatial proximity relationships between epithelial and immune subpopulations. Among epithelial subsets, only basal epithelial cells (Basal_Epi) cohabited niches with reparative fibroblast clusters, indicating Fib_6’s pro-regenerative signaling primarily targets basal epithelium. Fib_9’s spatial adjacency to hair follicle progenitors reinforced its papillary fibroblast identity (Supplementary Fig. 9A). Immune profiling revealed concentrated localization of Arg1^+^ macrophages, and Treg T cells within Fib_6-enriched niche_2 (Supplementary Fig. 9B-C), perfectly aligning with Fib_6-centric communication networks established in prior analyses.

### 3.6 EREG contributed to reparative fibroblast and regenerative pattern

To functionally validate regenerative niche signaling, recombinant PTN/EREG proteins (100 ng/mL) were administered peri-wound and intraperitoneally to young mice (6 week-old, n=5) from 14 to 21 dpw^54^, with healing outcomes assessed at 28 dpw (Fig. 6A). While control groups developed characteristic scar tissue, PTN treatment induced decreased collagen deposition without enhancing hair follicle regeneration (mean=2 vs control, p=0.5610) (Fig. 6B)^55^. In contrast, EREG significantly promoted de novo hair follicle formation (mean=9.6, p<0.01) with Ki-67+ proliferative HFs (Fig. 6C-D). ECM ultrastructure analysis revealed clustering differences between the PTN and normal groups. However, EREG-treated exhibited near-normal collagen architecture surrounded with fibrotic zones (Fig. 6E, Supplementary Fig. 10A-B) recapitulating aged regenerative patterns. In addition, CRABP1^+^/PRSS35^+^ reparative fibroblasts demonstrated preferential accumulation near EREG-induced HFs versus sparse papillary-layer distribution in PTN groups (Fig. 6F), establishing EREG’s superior capacity to mobilize regenerative subpopulations.

**Fig 6:**
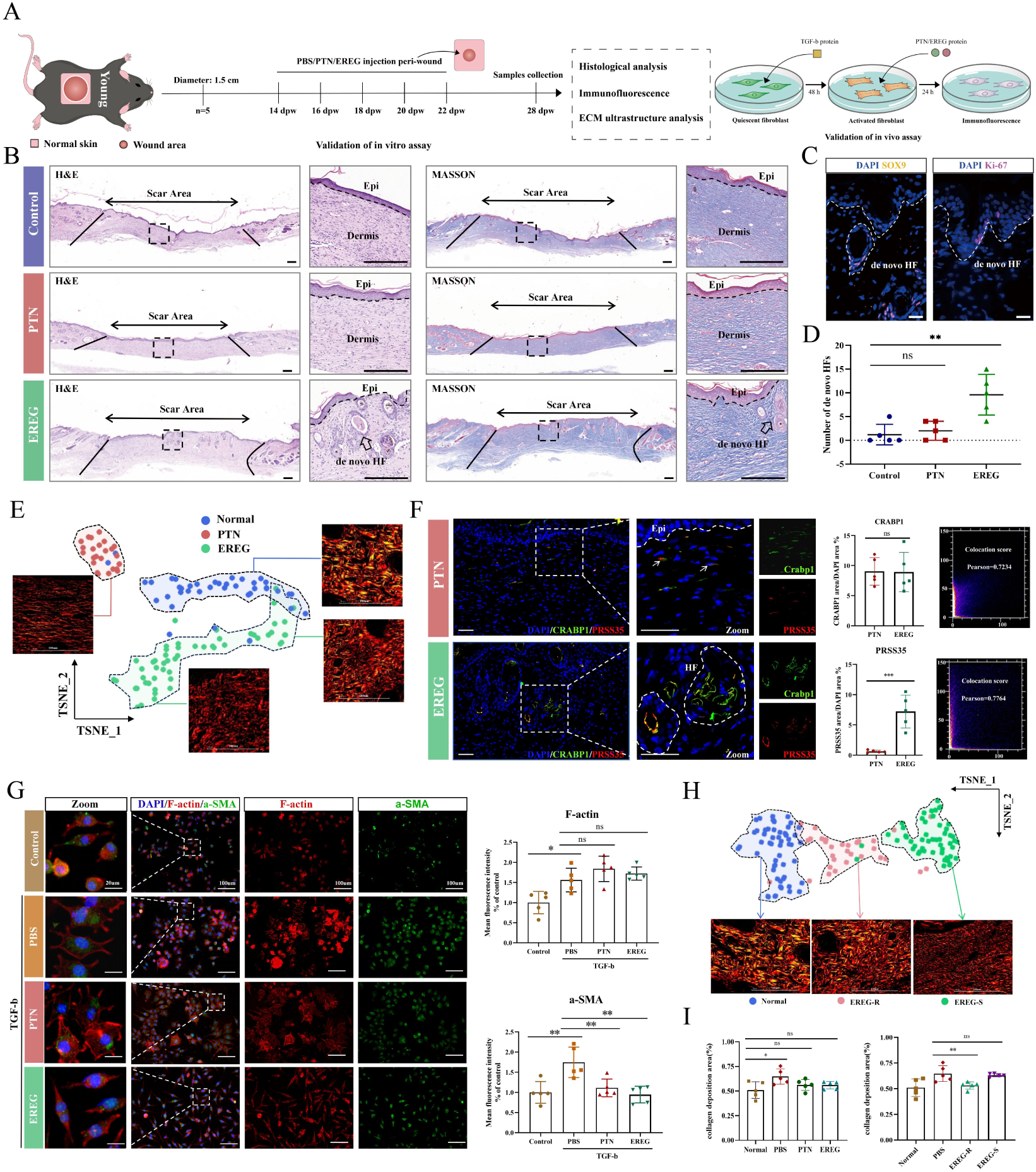
EREG contributed to reparative fibroblast and regenerative pattern. (A) Schematic diagram showing treatment design in vivo and vitro assays. (B)Hematoxylin-eosin (H&E) and Masson staining for scar tissues of control, PTN, and EREG treatment samples. Scale bar, 200 um. (C-D) Immunofluorescence staining for Sox9 (yellow) and Ki67 (purple) in the scar tissues of the EREG treatment group. Dotted lines signed morphology of de novo HF. Scale bar, 50um (C). Quantification analysis for numbers of de novo HFs (n=5 per group). Significances were utilized using bidirectional t-tests (D). (E) TSNE plots of unsupervised cluster analysis of ECM characteristics in each group with representative Sirius red staining images. Scale bar, 100um. (F) Immunofluorescence staining for CRABP1 (green, marker of papilla fibroblast), and PRSS35 (red, marker of reparative fibroblast) in the scar tissues. Scale bar, 50um(left). Quantification analysis for the ratio of marker protein area/DAPI area (n=5 per group). Significances were utilized using bidirectional t-tests(middle). Colocation plot showing colocalization of CRABP1/PRSS35 assessed by Pearson correlation coefficient(right). (G) Immunofluorescence staining for F-actin (red, cytoskeleton) and a-SMA (green, marker of fibroblast activation) in murine fibroblast lineage (L929 cells). Scale bar as shown in images. The difference in mean fluorescence intensity was utilized by bidirectional t-tests (n=5). (H-I) TSNE plots of unsupervised cluster analysis of ECM characteristics in the EREG group with representative Sirius red staining images. Scale bar, 100um(H). Quantification analysis for the ratio of collagen deposition area in scar tissues assessed by bidirectional t-tests (n=5). Abbreviation: HF, hair follicle. Epi, epidermis. EREG_R, EREG_regeneration area. EREG_S, EREG_scar area.a-SMA, a-smooth muscle actin. P value (*p < 0.05; **p < 0.01; ***p < 0.001), with nonsignificant (ns) determinations requiring p > 0.05.

Building on the hypothesis that reparative fibroblast subsets differentiate from myofibroblast, we examined PTN/EREG effects on fibroblast plasticity. TGF-β treatment of murine fibroblast lineage (L929) cells generated activated myofibroblasts^56^, exhibiting extended cytoskeletal F-actin filaments and upregulated α-SMA expression (p < 0.01). Following 24 h PTN/EREG treatment, both proteins reduced α-SMA levels (p < 0.01) without altering F-actin expression (p = 0.2115/PBS vs PTN; p = 0.5488/PBS vs EREG). Notably, EREG uniquely restored fibroblastic spindle morphology, contrasting with myofibroblast spreading morphology^39^, indicating its reduced fibrotic potential (Fig. 6G). Combined with the significant reduction of collagen deposition by EREG treatment, we demonstrate that EREG serves as a critical mediator enabling reparative fibroblasts to promote scarless wound healing (Fig. 6H-I).

### 3.7 Evaluation of the regenerative capacity of aged skin in small full-thickness wound

To assess age-related regenerative potential in small wound contexts, we established a 0.6 cm wound model^24,57^ and analyzed single-cell data from Vu Remy et al.’s 7 dpw small wound study^23^. Dimensionality reduction clustering resolved fibroblast, epithelial, and immune subpopulations, enabling comparative communication analysis (Fig. 7A). Results demonstrated concomitant upregulation of regenerative (WNT, BMP) and fibrotic (TGF-β) pathways in aged mice, while EGF signaling remained comparable (Fig. 7B), suggesting divergent regenerative mechanisms from large wounds. Longitudinal 28 dpw analysis of small wounds revealed non-significant hair follicle regeneration trends in aged mice (p=0.0711) (Fig. 7C-D). Collagen architecture analysis confirmed aged scars resembled young fibrotic structures (p=0.8652) (Fig. 7E-F, Supplementary Fig. 10C). IF detected abundant CRABP1+ fibroblasts but minimal PRSS35+ populations and EREG signaling in aged group (Fig. 7G-H), explaining incomplete regenerative outcomes despite latent capacity^13^ (Fig. 7I). Collectively, aged regenerative potential associates with reparative fibroblast subsets and EGF signaling, a program insufficiently activated in small wound contexts.

**Fig 7:**
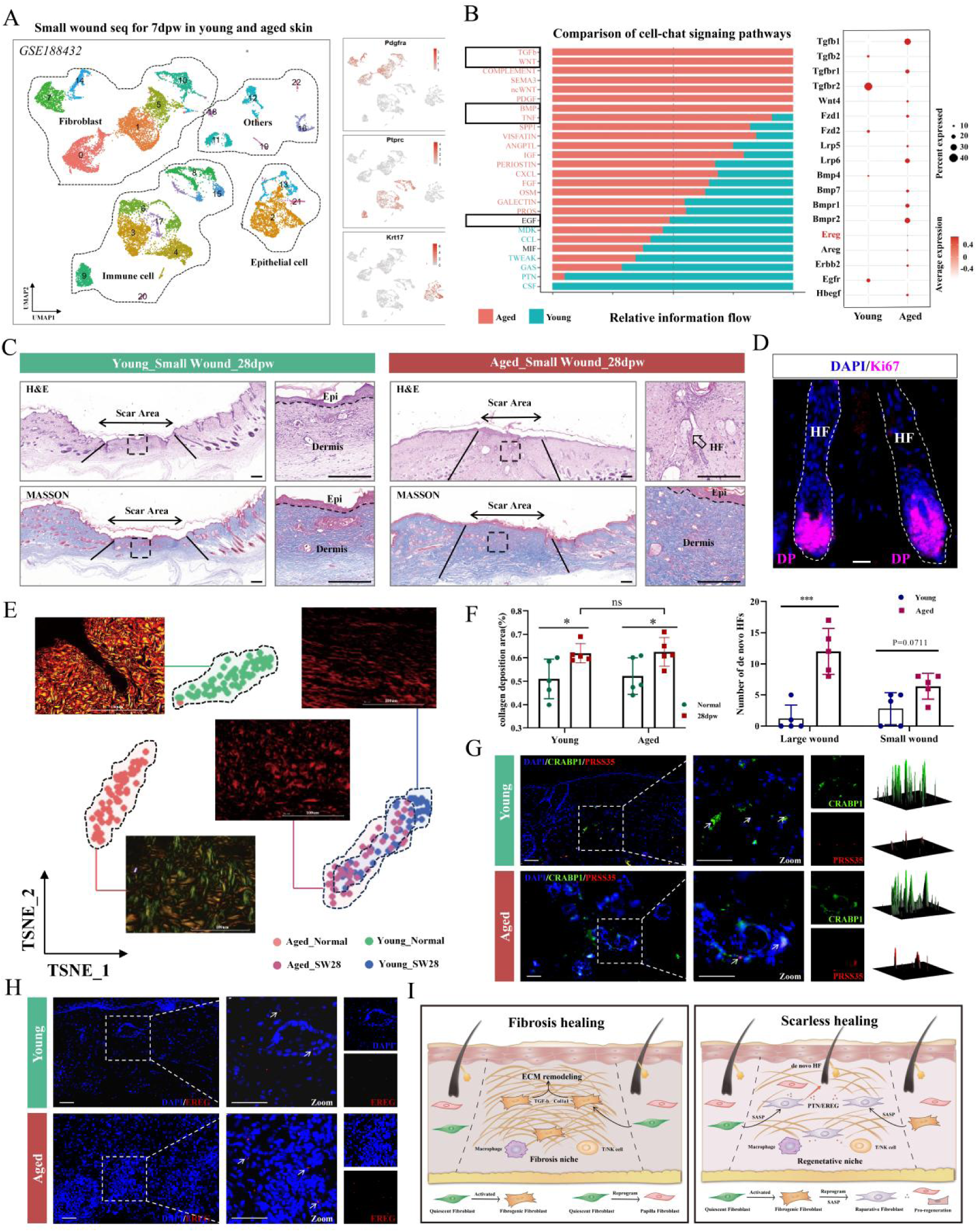
Evaluation of the regenerative capacity of aged skin in small full-thickness wound. (A) UMAP plots showing distribution and marker genes of major cell types from public small wound single-cell sequencing data(GSE188432). (B) Relative information flow showing up-regulated signaling between young and aged groups. Red, Aged specific. Blue, Young specific. The black signaling represents no differences(left). The dot plot showed the expression profile of relevant genes of characteristic signaling(right). (C) Hematoxylin-eosin (H&E) and Masson staining for scar tissues in young and aged small wound groups. Scale bar, 200 um. (D) Immunofluorescence staining for Ki67 (purple, marker of de novo HF) in the scar tissues of the aged group. Scale bar, 50um(upper). Quantification analysis for numbers of de novo HFs (n=5 per group). Significances were utilized using bidirectional t-tests(down). (E-F) TSNE plots of unsupervised cluster analysis of ECM characteristics in each group with representative Sirius red staining images. Scale bar, 100um(E). Quantification analysis for the ratio of collagen deposition area in scar tissues assessed by bidirectional t-tests(n=5)(F). (G) Immunofluorescence staining for CRABP1 (green, marker of papilla fibroblast), and PRSS35 (red, marker of reparative fibroblast) in the scar tissues(left). Surface plot showing relative fluorescence intensity of marker protein(right). Scale bar, 50 um. (H) Immunofluorescence staining for EREG(green) in the scar tissues between young and aged groups. Scale bar, 50um. Surface plot showing relative fluorescence intensity of EREG. (I) Schematic diagram showing pivotal distinct signaling and cell types in fibrotic and regenerative healing. Abbreviation: HF, hair follicle. Epi, epidermis.DP, dermal papilla. SW, small wound. Statistical thresholds followed p = 0.05 convention (*p < 0.05; **p < 0.01; ***p < 0.001; ****p < 0.0001), with nonsignificant (ns) determinations requiring p > 0.05.

## Discussion

By integrating single-cell sequencing, spatial transcriptomics, and functional validation, we revealed that aged mice exhibited more pronounced regenerative capacity compared to young mice after large wound healing, characterized by increased de nove HF numbers and collagen fiber features closer to normal skin. Single-cell sequencing identified a reparative fibroblast subpopulation (Prss35+Fib) enriched in the aged group, which promotes regeneration through communication with epithelial cells, macrophages, and T cells via PTN and EREG signaling. Spatial transcriptomics validated this communication pattern by elucidating cell proximity and locating the regenerative niche in the upper dermis. Finally, EREG treatment significantly enhanced regenerative outcomes in young mice and suppressed fibrotic markers of myofibroblast. While the small wound model of aged mice, lacking the reparative fibroblast(Prss35+Fib) and EREG signaling, failed to achieve scarless healing.

While our findings demonstrate aged skin exhibits regenerative healing in large full-thickness wounds, this capacity remains dormant in small wound models – implicating age-independent factors like mechanical stress in regenerative potential activation. Prior work in wound-induced HF regeneration models reveals scar-center regenerative areas with permissive stiffness ranges (5-15 kPa) versus peripheral scar zones exhibiting biomechanical restriction (>15 kPa), a stratification mediated by collagen composition heterogeneity across scar regions^58,59^. Notably, aged large wounds and EREG-treated young scars exhibit collagen architectures approximating unwounded skin (Fig. 7F-G), a structural similarity potentially enabling permissive stiffness for HF regeneration. In contrast, small wounds in aged and young mice display analogous type I collagen alignment patterns distinct from physiologic dermal organization, likely explaining their shared regenerative failure. In addition, mechanotransductive pathways like En1-mediated signaling critically regulates fibroblast fate determination in scars, suggesting a molecular framework for biomechanical control of regeneration^20,59^.

Notably, reparative fibroblasts (Prss35*^+^*) were consistently observed within regenerated hair follicles across aged wound models (regardless of injury scale), suggesting Prss35+ fibroblast abundance may dictate healing modality transitions. Intriguingly, reindeer antler skin (a natural regenerative model) harbors analogous *Prss35+* fibroblasts co-expressing PTN/MDK, further supporting their conserved regenerative role^2^. However, cross-species extrapolation to aged human skin remains unverified, as existing public datasets solely profile unwounded aged skin, not post-healing scar tissue^60,61^. The studies of human wound healing revealed that EGFR signaling critically regulates re-epithelialization, mirroring Prss35+ fibroblasts’ secretory profile^62^. Thus, interrogating Prss35 overexpression in skin organoids for HF neogenesis, or analyzing aged patient scar specimens for Prss35+ fibroblast correlates, could clarify the cross-species conservation of regenetation potential^63^.

Building on the essential roles of cellular senescence and reprogramming in tissue regeneration, we hypothesize reparative subpopulation reprogramming is driven by SASP (EREG)^64^. Prior studies indicate aging enhances rather than impedes reprogramming efficacy, and profound epigenetic remodeling may render cells more amenable to stem-like transcriptional reprogramming^45,65^. For instance, fibroblasts of aged mice exhibit heightened reprogramming capacity into pluripotent stem cells under inflammatory cues (e.g., IL-6) compared to young counterparts^66^. Future investigations should delineate the relationship between SASP and fibroblast reprogramming, such as EREG/EGFR downstream effectors, and potential synergistic interactions with pathways like the Wnt signaling pathway^54,67^.

Collectively, our findings reframe aging-regeneration relationships: aged skin establishes reparative fibroblast-enriched niches post-injury, orchestrating HF regeneration and collagen normalization, albeit amidst persistent fibrotic regions. Therapeutic exploitation through niche signal modulation (e.g., PTN/MDK crosstalk), and SASP-factor (IL-6/EREG) biomaterial delivery systems, may achieve complete scarless regeneration.

## Method

### 2.1 Animals and excisional wound model

The study was approved by the Animal Research Committee of Sichuan University. Male C57BL/6 mice were purchased from Chengdu Dossy Experimental Animals Co., Ltd (China) and housed in appropriate environments. The excisional wound model described below was according to established protocols^19,24^. A full-thickness circular wound(diameter=1.5 cm/0.6cm) was created in the middle of dorsal skin and stented open by a silicone ring, which was fixed with sutures and maintained until re-epithelialization was complete. Aseptic dressings(3M) were applied over the silicone ring around the wound and changed at regular intervals. At the indicated day post-wound (dpw), the wound samples were harvested under anesthesia. Normal skin samples were harvested during the creation of the wound model.

### 2.2 Histology and Immunofluorescence

After fixation in 4% paraformaldehyde for 24 hours, the samples were dehydrated, cleared, and subsequently embedded in paraffin at a 45-degree angle. Trim the embedded samples until reaching the scar area devoid of panniculus carnosus. Sections were performed with H&E staining to observe the wound healing process, while Masson and Sirius red staining were performed to assess the deposition of collagen fiber. Immunofluorescence staining was performed following established protocols. Ki-67( ) and SOX9( ) were utilized for observation of hair follicle, while CRABP1(12588-1-AP, proteintech), a-SMA(67735-1-Ig, proteintech), PRSS35(GTX123037, GeneTex), PTN(HA722055, HUABIO), and EREG(ER65835, HUABIO) were employed to assess the fibroblast subcluster.

### 2.3 ECM ultrastructural analysis

For quantitative analysis of the ECM ultrastructure, the samples were divided into six groups(Young: normal, 21dpw, 28dpw; Aged: normal, 21dpw, 28dpw). Sections performed by Sirius red staining(5 samples per group, 2 sections per sample, 6 sites per section) were imaged using a polarized light microscope at 40x magnification. Stored images were normalized and color deconvoluted per the established algorithm to isolate individual colors within RGB channels in Matlab R2023b^68^. Isolated red images represent mature fibers and green images represent immature fibers, as explained in previous reports^19,20^. Afterward, the noise of isolated images was removed with adaptive filtering using the wiener2 function, which preserves regions with high variance(edge of fibers). The filter images(Grayscale images) were then binarized using the im2bw function and selected for the diamond structuring element using the imerode function to obtain Binary images. At last, Skeletonized images were harvested by skeletonizing the Binary images using the bwmorph function. Parameters of the digitized images(as listed in Table S1) were measured using the regionprops function to obtain quantified properties of fibers (ECM ultrastructure). Dimension reduction and visualization of the quantified result were performed with the t-distributed stochastic neighbor embedding (t-SNE, v0.1) package in R(4.1.2). Matlab pipelines described above are available at: https://github.com/shamikmascharak/Mascharak-et-al-ENF.

### 2.4 Bulk RNA sequencing

Three replicate samples of scar tissue per group were processed for bulk RNA-seq. Total RNA was extracted using TRIzol(Invitrogen), assessed for quality, and quantified (cat. 61006; Thermo). Poly-A mRNA was enriched, fragmented, and reverse-transcribed into cDNA (SuperScript kit). Libraries were constructed via end-repair, adapter ligation, PCR amplification (Phusion DNA polymerase), and size selection (300 bp fragments) following Illumina protocols. Sequencing was performed on Illumina NovaSeq 6000 (2×150 bp reads). Raw reads were trimmed, aligned to the reference genome, and quantified. Differential expression gene analysis (DESeq2; |log2FC|>1, p<0.05) and functional enrichment (ClusterProfiler; GO/GSEA, p<0.05) were conducted subsequently.

### 2.5 single-cell RNA sequencing (scRNA-seq) Tissue dissociation and Sequence

Following two rinses in cold PBS, the wound samples were cut into 0.5mm^3^ pieces and digested with enzymatic hydrolysate for 60 minutes at 37 ℃. After repeated filtration, centrifugation, and suspension, red blood cells (MACS Red Blood Cell Lysis, 130-094-183) and dead cells (MACS Dead Cell Removal Kit, 130-090-101) were removed from the cell suspension. Then single cell of the prepared cell suspensions was labeled and carried out reverse transcription to construct a cDNA library using the 10 × Genomics Chromium Next GEM Single Cell 3 ʹ Reagent Kits v3.1(1000268). The constructed library was sequenced using the Illumina NOVA 6000 PE150 platform.

#### Data processed and cell type annotation

The raw sequencing data was demultiplexed and aligned using Cell Ranger software, and subsequent analyses were performed using R (v4.1.2) and Seruat packages (v4.1.1). Quality control was applied to identify low-quality cells, defined by mitochondrial gene expression ratios of less than 10% and minimum gene expression numbers of more than 200. Data normalization along with batch effects removal was performed to generate a comparable gene expression matrix. The first 2000 highly variable genes were identified using the FindVariableFeatures function and utilized for principal component analysis. Dimension reduction was achieved based on the first 20 principal components using the RunUMAP function. Subsequently, the FindNeighbors and FindClusters functions were used for clustering the data, while the marker genes of clustering results were calculated using the FindAllMarkers function, facilitating the annotation of distinct cell types. Gene enrichment analysis was performed using GO.db(v3.14.0) and ClusterProfiler(v4.2.2) packages.

#### Pseudotime analysis

Pseudotime analysis was performed using the Monocle2 package (v2.9.0) to infer the differentiation trajectories of specified cells: Initially, the Seurat object was converted to the CellDataSet object using the ImportCDS function, and ordered genes (q-value < 0.01) was selected by DifferentialGeneTest function. Then the reduceDimension function was employed to perform dimension-reduced clustering to build spanning trees, which facilitates trajectory inference based on the orderCells function. Additionally, the Slingshot package(v2.10.0) was utilized for cross-validation of the predicted trajectory above.

#### Cytotrace analysis

With the cytotrace package (v0.3.3), the predicted order of fibroblast subclusters was assessed. The predicted order at 0 represents a high differentiation state and low development potential, while that at 1 represents the contrary. In brief, cytotrace employs a straightforward and reliable determinant of developmental potential — the number of detectable expressed genes per cell to predict cell states. As cellular differentiation progresses, the number of expressed genes and developmental potential decreases^37^.

#### RNA velocity analysis

RNA velocity analysis solves the complete transcription of splicing dynamics using a likelihood-based kinetic model with the scvelo package^38^. Its principle lies in inferring gene-specific velocity of transcription, splicing, and degradation, to reconstruct the position of each cell along potential differentiation trajectories, and to detect putative driving genes. By projecting the velocity vectors onto UMAP embedding, the differentiation trajectory of individual cell can be predicted based on transition probabilities.

#### Cellchat receptor-ligand analysis

With the CellChat package(V1.6.1), cell-cell communication analysis was conducted to infer the receptor-ligand networks among cell subclusters. Initially, the Seurat object was converted to a CellChat object using the creaCellChat function, with the incorporation of the ligand-receptor interaction database using CellChatDB.mouse. Then the gene expression data was extracted using cellchat@data.signaling and utilized for identifying overexpressed genes and ligand-receptor pairs. Subsequently, the communication probabilities were computed based on the ligand-receptor pairs and P-values.

### 2.6 10x Visium CyAssist Spatial transcriptomics (ST) Tissue preparation and Sequence

Freshly collected wound samples were trimmed until reaching the scar area and embedded in OCT. After methanol fixation, H&E staining, and imaging, appropriate sections were selected for destaining and reverse transcription for cDNA synthesis. The gene expression map of the section was anchored, through probe hybridizing, releasing, and transferring the cDNA to the 10xVisium CytAssist slide(10xGenomics, CG000495). Then the cDNA library was constructed with the Visium CytAssist Spatial Gene Expression for FFPE kit(PN-1000521,6.5mm) and sequenced using the PE-150 model.

#### Spot deconvolution and annotation

The raw sequencing data was assessed and normalized using the Space Ranger software and the SCTtransform function of the Seurat package in R(variable features=3000). As described in scRNA-seq above, clusters were identified with the FindNeighbors function(dims=1:30), the FindClusters function (resolution=0.4), and the RunUMAP function. To inference cell types of the individual spot from the sequencing data, the RCTD algorithm of the spacexr package(v2.2.1) was performed to integrate signatures of scRNA-seq and ST data. In brief, RCTD deconvolutes the gene expression matrix of ST data following signatures of scRNA-seq reference to obtain annotation of cell types per spot.

#### Cell type niche definition

Niche analysis was performed to identify distinct niches in specific spatial regions, defined by spots with proximity cell composition. Specifically, the niche represents distinctive cell types and the spatial and functional dependencies among them. Initially, the cell type composition of spots was converted into an isometric log ratio (ILR) and carried out dimension reduction using the uwot package (v0.1.14). Then the niche summary matrix generated was hierarchically clustered using the hclust function. At last, the Wilcoxon test was performed to assess the significance of selected cell types (p-value <0.05).

#### cell-cell colocalization

With the mistyR package(v1.2.1), the importance of the spatial dependence of selected cell types was assessed, which presents the result of cell-cell colocalization. Initially, two views of the spatial pattern were defined (intra view describes predicted result within one spot, intra = Null; para view describes result of peripheral neighbors, para =5). In accordance with deconvolution per spot, the importance of target cell types in individual view was calculated, collected, and aggregated using misty workflow in the source of misty_utilities.R. The importance score of the predictor was interpreted as spatial dependence with target cell type^52^.

#### Cell-cell Ligand-receptor interactions

With the stLearn package(v0.4.12) in the Python environment, ligand-receptor interactions within spatial slides were identified and predicted. The raw sequencing data was converted into an anndata object in jupyterlab platform, with incorporation of the interaction database using connectomeDB2020_list, mouse. Then CCI analysis (min_spots=20, n_pairs=10000) was employed for detecting ligand-receptor expression in spots, with a permutation test to calculate significance. Afterward, based on the combination of the ligand-receptor results with the spot deconvolution results, the cell-cell communication network was predicted (min_spots=5, cell_prop_cutoff=0.1, n_prems=1000).

### 2.7 Quantitation and statistical analysis

All quantitative comparisons leveraged GraphPad Prism 8.0 and image J with study design-appropriate tests: paired comparisons utilized bidirectional t-tests, whereas multigroup comparisons against a single control group employed one-way ANOVA with Dunnett’s post hoc analysis. Spatial colocalization was resolved through Pearson correlation-based quantification. Statistical thresholds followed p = 0.05 convention.

## Data availability

The sequencing data of this study are available.

## Code availability

R code is available from the corresponding author upon request.

## Acknowledgements

This study was supported by National Natural Science Foundation of China (NSFC, 81970966) and Natural Science Foundation of Sichuan Province (2023NSFSC0574). We thank OE Biotech Co., Ltd (Shanghai, China) for providing RNA sequence, single cell RNA sequence and spatial RNA sequence, kunyue meng and lanying wang and hongyan liu for assistance with bioinformatics analysis.

## Contributions

D.Y.W., Z.Q.W., and Y.Y. designed the study, conducted experiments, and analyzed the data. F.W.B. and H.Y.L. assisted in animal experiments. J.N.H. assisted in schematic drawing. F.Z., T.C., F.T.F., Z.Y.W., and J.H.Z assisted in data discussion. D.Y.W., Z.Q.W., and Y.Y. prepared the manuscript with the guidance of L.X., Y.M., and Y.Y.W.

These authors contributed equally to this study: Dongyang Wang, Zhanqi Wang, Yang Yang

## Reference

1. Clark, R.A.F. (2021). To Scar or Not to Scar. The New England journal of medicine 385, 469–471. 10.1056/NEJMcibr2107204.

2. Sinha, S., Sparks, H.D., Labit, E., Robbins, H.N., Gowing, K., Jaffer, A., Kutluberk, E., Arora, R., Raredon, M.S.B., Cao, L., et al. (2022). Fibroblast inflammatory priming determines regenerative versus fibrotic skin repair in reindeer. Cell 185, 4717–4736.e4725. 10.1016/j.cell.2022.11.004.

3. Seifert, A.W., Kiama, S.G., Seifert, M.G., Goheen, J.R., Palmer, T.M., and Maden, M. (2012). Skin shedding and tissue regeneration in African spiny mice (Acomys). Nature 489, 561–565. 10.1038/nature11499.

4. Willyard, C. (2018). Unlocking the secrets of scar-free skin healing. Nature 563, S86–s88. 10.1038/d41586-018-07430-w.

5. Zhao, X., Kwan, J.Y.Y., Yip, K., Liu, P.P., and Liu, F.F. (2020). Targeting metabolic dysregulation for fibrosis therapy. Nat Rev Drug Discov 19, 57–75. 10.1038/s41573-019-0040-5.

6. Global burden of 369 diseases and injuries in 204 countries and territories, 1990-2019: a systematic analysis for the Global Burden of Disease Study 2019. (2020). Lancet (London, England) 396, 1204–1222. 10.1016/s0140-6736(20)30925-9.

7. Mascharak, S., desJardins-Park, H.E., and Longaker, M.T. (2020). Fibroblast Heterogeneity in Wound Healing: Hurdles to Clinical Translation. Trends Mol Med 26, 1101–1106. 10.1016/j.molmed.2020.07.008.

8. Abbasi, S., Sinha, S., Labit, E., Rosin, N.L., Yoon, G., Rahmani, W., Jaffer, A., Sharma, N., Hagner, A., Shah, P., et al. (2020). Distinct Regulatory Programs Control the Latent Regenerative Potential of Dermal Fibroblasts during Wound Healing. Cell Stem Cell 27, 396–412.e396. 10.1016/j.stem.2020.07.008.

9. Capolupo, L., Khven, I., Lederer, A.R., Mazzeo, L., Glousker, G., Ho, S., Russo, F., Montoya, J.P., Bhandari, D.R., Bowman, A.P., et al. (2022). Sphingolipids control dermal fibroblast heterogeneity. Science 376, eabh1623. 10.1126/science.abh1623.

10. Yin, J.L., Wu, Y., Yuan, Z.W., Gao, X.H., and Chen, H.D. (2020). Advances in scarless foetal wound healing and prospects for scar reduction in adults. Cell proliferation 53, e12916. 10.1111/cpr.12916.

11. Cai, Y., Xiong, M., Xin, Z., Liu, C., Ren, J., Yang, X., Lei, J., Li, W., Liu, F., Chu, Q., et al. (2023). Decoding aging-dependent regenerative decline across tissues at single-cell resolution. Cell Stem Cell 30, 1674–1691.e1678. 10.1016/j.stem.2023.09.014.

12. Chen, S.D., Chu, C.Y., Wang, C.B., Yang, Y., Xu, Z.Y., Qu, Y.L., and Man, Y. (2024). Integrated-omics profiling unveils the disparities of host defense to ECM scaffolds during wound healing in aged individuals. Biomaterials 311, 122685. 10.1016/j.biomaterials.2024.122685.

13. Nishiguchi, M.A., Spencer, C.A., Leung, D.H., and Leung, T.H. (2018). Aging Suppresses Skin-Derived Circulating SDF1 to Promote Full-Thickness Tissue Regeneration. Cell Rep 24, 3383–3392.e3385. 10.1016/j.celrep.2018.08.054.

14. de Magalhães, J.P. (2024). Cellular senescence in normal physiology. Science 384, 1300–1301. 10.1126/science.adj7050.

15. Da Silva-Álvarez, S., Guerra-Varela, J., Sobrido-Cameán, D., Quelle, A., Barreiro-Iglesias, A., Sánchez, L., and Collado, M. (2020). Cell senescence contributes to tissue regeneration in zebrafish. Aging cell 19, e13052. 10.1111/acel.13052.

16. Salinas-Saavedra, M., Febrimarsa, Krasovec, G., Horkan, H.R., Baxevanis, A.D., and Frank, U. (2023). Senescence-induced cellular reprogramming drives cnidarian whole-body regeneration. Cell Rep 42, 112687. 10.1016/j.celrep.2023.112687.

17. Walters, H.E., Troyanovskiy, K.E., Graf, A.M., and Yun, M.H. (2023). Senescent cells enhance newt limb regeneration by promoting muscle dedifferentiation. Aging cell 22, e13826. 10.1111/acel.13826.

18. Plikus, M.V., Wang, X., Sinha, S., Forte, E., Thompson, S.M., Herzog, E.L., Driskell, R.R., Rosenthal, N., Biernaskie, J., and Horsley, V. (2021). Fibroblasts: Origins, definitions, and functions in health and disease. Cell 184, 3852–3872. 10.1016/j.cell.2021.06.024.

19. Mascharak, S., Talbott, H.E., Januszyk, M., Griffin, M., Chen, K., Davitt, M.F., Demeter, J., Henn, D., Bonham, C.A., Foster, D.S., et al. (2022). Multi-omic analysis reveals divergent molecular events in scarring and regenerative wound healing. Cell Stem Cell 29, 315–327.e316. 10.1016/j.stem.2021.12.011.

20. Mascharak, S., desJardins-Park, H.E., Davitt, M.F., Griffin, M., Borrelli, M.R., Moore, A.L., Chen, K., Duoto, B., Chinta, M., Foster, D.S., et al. (2021). Preventing Engrailed-1 activation in fibroblasts yields wound regeneration without scarring. Science 372. 10.1126/science.aba2374.

21. Plikus, M.V., Guerrero-Juarez, C.F., Ito, M., Li, Y.R., Dedhia, P.H., Zheng, Y., Shao, M., Gay, D.L., Ramos, R., Hsi, T.C., et al. (2017). Regeneration of fat cells from myofibroblasts during wound healing. Science 355, 748–752. 10.1126/science.aai8792.

22. Hinz, B., Phan, S.H., Thannickal, V.J., Prunotto, M., Desmoulière, A., Varga, J., De Wever, O., Mareel, M., and Gabbiani, G. (2012). Recent developments in myofibroblast biology: paradigms for connective tissue remodeling. Am J Pathol 180, 1340–1355. 10.1016/j.ajpath.2012.02.004.

23. Vu, R., Jin, S., Sun, P., Haensel, D., Nguyen, Q.H., Dragan, M., Kessenbrock, K., Nie, Q., and Dai, X. (2022). Wound healing in aged skin exhibits systems-level alterations in cellular composition and cell-cell communication. Cell Rep 40, 111155. 10.1016/j.celrep.2022.111155.

24. Yang, Y., Chu, C., Liu, L., Wang, C., Hu, C., Rung, S., Man, Y., and Qu, Y. (2023). Tracing immune cells around biomaterials with spatial anchors during large-scale wound regeneration. Nat Commun 14, 5995. 10.1038/s41467-023-41608-9.

25. Peña, O.A., and Martin, P. (2024). Cellular and molecular mechanisms of skin wound healing. Nat Rev Mol Cell Biol 25, 599–616. 10.1038/s41580-024-00715-1.

26. Li, H., Wang, Z., Zhou, F., Zhang, G., Feng, X., Xiong, Y., and Wu, Y. (2023). Sustained activation of NLRP3 inflammasome contributes to delayed wound healing in aged mice. International immunopharmacology 116, 109828. 10.1016/j.intimp.2023.109828.

27. Nanba, D., Toki, F., Asakawa, K., Matsumura, H., Shiraishi, K., Sayama, K., Matsuzaki, K., Toki, H., and Nishimura, E.K. (2021). EGFR-mediated epidermal stem cell motility drives skin regeneration through COL17A1 proteolysis. J Cell Biol 220. 10.1083/jcb.202012073.

28. Li, X., An, T., Yang, Y., Xu, Z., Chen, S., Yi, Z., Deng, C., Zhou, F., Man, Y., and Hu, C. (2024). TLR9 activation in large wound induces tissue repair and hair follicle regeneration via γδ T cells. Cell Death Dis 15, 598. 10.1038/s41419-024-06994-y.

29. Sun, Z., Chen, Z., Yin, M., Wu, X., Guo, B., Cheng, X., Quan, R., Sun, Y., Zhang, Q., Fan, Y., et al. (2024). Harnessing developmental dynamics of spinal cord extracellular matrix improves regenerative potential of spinal cord organoids. Cell Stem Cell 31, 772–787.e711. 10.1016/j.stem.2024.03.007.

30. Deguchi, E., Lin, S., Hirayama, D., Matsuda, K., Tanave, A., Sumiyama, K., Tsukiji, S., Otani, T., Furuse, M., Sorkin, A., et al. (2024). Low-affinity ligands of the epidermal growth factor receptor are long-range signal transmitters in collective cell migration of epithelial cells. Cell Rep 43, 114986. 10.1016/j.celrep.2024.114986.

31. Schmidt, I.M., Colona, M.R., Kestenbaum, B.R., Alexopoulos, L.G., Palsson, R., Srivastava, A., Liu, J., Stillman, I.E., Rennke, H.G., Vaidya, V.S., et al. (2021). Cadherin-11, Sparc-related modular calcium binding protein-2, and Pigment epithelium-derived factor are promising non-invasive biomarkers of kidney fibrosis. Kidney Int 100, 672–683. 10.1016/j.kint.2021.04.037.

32. McKenney, C., Lendner, Y., Guerrero Zuniga, A., Sinha, N., Veresko, B., Aikin, T.J., and Regot, S. (2024). CDK4/6 activity is required during G(2) arrest to prevent stress-induced endoreplication. Science 384, eadi2421. 10.1126/science.adi2421.

33. Gopee, N.H., Winheim, E., Olabi, B., Admane, C., Foster, A.R., Huang, N., Botting, R.A., Torabi, F., Sumanaweera, D., Le, A.P., et al. (2024). A prenatal skin atlas reveals immune regulation of human skin morphogenesis. Nature 635, 679–689. 10.1038/s41586-024-08002-x.

34. Kanaan, R., Medlej-Hashim, M., Jounblat, R., Pilecki, B., and Sorensen, G.L. (2022). Microfibrillar-associated protein 4 in health and disease. Matrix Biol 111, 1–25. 10.1016/j.matbio.2022.05.008.

35. Lim, C.H., Sun, Q., Ratti, K., Lee, S.H., Zheng, Y., Takeo, M., Lee, W., Rabbani, P., Plikus, M.V., Cain, J.E., et al. (2018). Hedgehog stimulates hair follicle neogenesis by creating inductive dermis during murine skin wound healing. Nat Commun 9, 4903. 10.1038/s41467-018-07142-9.

36. Li, B., Zhang, W., Guo, C., Xu, H., Li, L., Fang, M., Hu, Y., Zhang, X., Yao, X., Tang, M., et al. (2022). Benchmarking spatial and single-cell transcriptomics integration methods for transcript distribution prediction and cell type deconvolution. Nat Methods 19, 662–670. 10.1038/s41592-022-01480-9.

37. Gulati, G.S., Sikandar, S.S., Wesche, D.J., Manjunath, A., Bharadwaj, A., Berger, M.J., Ilagan, F., Kuo, A.H., Hsieh, R.W., Cai, S., et al. (2020). Single-cell transcriptional diversity is a hallmark of developmental potential. Science 367, 405–411. 10.1126/science.aax0249.

38. Bergen, V., Lange, M., Peidli, S., Wolf, F.A., and Theis, F.J. (2020). Generalizing RNA velocity to transient cell states through dynamical modeling. Nat Biotechnol 38, 1408–1414. 10.1038/s41587-020-0591-3.

39. Younesi, F.S., Miller, A.E., Barker, T.H., Rossi, F.M.V., and Hinz, B. (2024). Fibroblast and myofibroblast activation in normal tissue repair and fibrosis. Nat Rev Mol Cell Biol 25, 617–638. 10.1038/s41580-024-00716-0.

40. Zhang, J., Wang, H., Chen, H., Li, H., Xu, P., Liu, B., Zhang, Q., Lv, C., and Song, X. (2022). ATF3 -activated accelerating effect of LINC00941/lncIAPF on fibroblast-to-myofibroblast differentiation by blocking autophagy depending on ELAVL1/HuR in pulmonary fibrosis. Autophagy 18, 2636–2655. 10.1080/15548627.2022.2046448.

41. Wilsker, D., Patsialou, A., Dallas, P.B., and Moran, E. (2002). ARID proteins: a diverse family of DNA binding proteins implicated in the control of cell growth, differentiation, and development. Cell Growth Differ 13, 95–106.

42. Subbaiah, K.C.V., Wu, J., Tang, W.H.W., and Yao, P. (2022). FAM114A1 influences cardiac pathological remodeling by regulating angiotensin II signaling. JCI insight 7. 10.1172/jci.insight.152783.

43. Gerlach, J.P., Jordens, I., Tauriello, D.V.F., van ‘t Land-Kuper, I., Bugter, J.M., Noordstra, I., van der Kooij, J., Low, T.Y., Pimentel-Muiños, F.X., Xanthakis, D., et al. (2018). TMEM59 potentiates Wnt signaling by promoting signalosome formation. Proc Natl Acad Sci U S A 115, E3996–e4005. 10.1073/pnas.1721321115.

44. Lucas, V., Cavadas, C., and Aveleira, C.A. (2023). Cellular Senescence: From Mechanisms to Current Biomarkers and Senotherapies. Pharmacol Rev 75, 675–713. 10.1124/pharmrev.122.000622.

45. Reimann, M., Lee, S., and Schmitt, C.A. (2024). Cellular senescence: Neither irreversible nor reversible. J Exp Med 221. 10.1084/jem.20232136.

46. Reyes, N.S., Krasilnikov, M., Allen, N.C., Lee, J.Y., Hyams, B., Zhou, M., Ravishankar, S., Cassandras, M., Wang, C., Khan, I., et al. (2022). Sentinel p16(INK4a+) cells in the basement membrane form a reparative niche in the lung. Science 378, 192–201. 10.1126/science.abf3326.

47. Takeo, M., Toyoshima, K.E., Fujimoto, R., Iga, T., Takase, M., Ogawa, M., and Tsuji, T. (2023). Cyclical dermal micro-niche switching governs the morphological infradian rhythm of mouse zigzag hair. Nat Commun 14, 4478. 10.1038/s41467-023-39605-z.

48. Choi, N., Kim, W.S., Oh, S.H., and Sung, J.H. (2020). Epiregulin promotes hair growth via EGFR-medicated epidermal and ErbB4-mediated dermal stimulation. Cell proliferation 53, e12881. 10.1111/cpr.12881.

49. Ng, M.T.H., Borst, R., Gacaferi, H., Davidson, S., Ackerman, J.E., Johnson, P.A., Machado, C.C., Reekie, I., Attar, M., Windell, D., et al. (2024). A single cell atlas of frozen shoulder capsule identifies features associated with inflammatory fibrosis resolution. Nat Commun 15, 1394. 10.1038/s41467-024-45341-9.

50. Konieczny, P., Xing, Y., Sidhu, I., Subudhi, I., Mansfield, K.P., Hsieh, B., Biancur, D.E., Larsen, S.B., Cammer, M., Li, D., et al. (2022). Interleukin-17 governs hypoxic adaptation of injured epithelium. Science 377, eabg9302. 10.1126/science.abg9302.

51. Gao, S., Shi, Q., Zhang, Y., Liang, G., Kang, Z., Huang, B., Ma, D., Wang, L., Jiao, J., Fang, X., et al. (2022). Identification of HSC/MPP expansion units in fetal liver by single-cell spatiotemporal transcriptomics. Cell Res 32, 38–53. 10.1038/s41422-021-00540-7.

52. Kuppe, C., Ramirez Flores, R.O., Li, Z., Hayat, S., Levinson, R.T., Liao, X., Hannani, M.T., Tanevski, J., Wünnemann, F., Nagai, J.S., et al. (2022). Spatial multi-omic map of human myocardial infarction. Nature 608, 766–777. 10.1038/s41586-022-05060-x.

53. Sänger, C.S., Cernakova, M., Wietecha, M.S., Garau Paganella, L., Labouesse, C., Dudaryeva, O.Y., Roubaty, C., Stumpe, M., Mazza, E., Tibbitt, M.W., et al. (2023). Serine protease 35 regulates the fibroblast matrisome in response to hyperosmotic stress. Sci Adv 9, eadh9219. 10.1126/sciadv.adh9219.

54. Lemmetyinen, T.T., Viitala, E.W., Wartiovaara, L., Kaprio, T., Hagström, J., Haglund, C., Katajisto, P., Wang, T.C., Domènech-Moreno, E., and Ollila, S. (2023). Fibroblast-derived EGF ligand neuregulin 1 induces fetal-like reprogramming of the intestinal epithelium without supporting tumorigenic growth. Disease models & mechanisms 16. 10.1242/dmm.049692.

55. Lamprou, M., Kastana, P., Kofina, F., Tzoupis, Η., Barmpoutsi, S., Sajib, M.S., Koutsioumpa, M., Poimenidi, E., Zompra, A.A., Tassopoulos, D., et al. (2020). Pleiotrophin selectively binds to vascular endothelial growth factor receptor 2 and inhibits or stimulates cell migration depending on α ( ν ) β (3) integrin expression. Angiogenesis 23, 621–636. 10.1007/s10456-020-09733-x.

56. Lodyga, M., and Hinz, B. (2020). TGF-β1 - A truly transforming growth factor in fibrosis and immunity. Semin Cell Dev Biol 101, 123–139. 10.1016/j.semcdb.2019.12.010.

57. Phan, Q.M., Sinha, S., Biernaskie, J., and Driskell, R.R. (2021). Single-cell transcriptomic analysis of small and large wounds reveals the distinct spatial organization of regenerative fibroblasts. Experimental dermatology 30, 92–101. 10.1111/exd.14244.

58. Harn, H.I., Wang, S.P., Lai, Y.C., Van Handel, B., Liang, Y.C., Tsai, S., Schiessl, I.M., Sarkar, A., Xi, H., Hughes, M., et al. (2021). Symmetry breaking of tissue mechanics in wound induced hair follicle regeneration of laboratory and spiny mice. Nat Commun 12, 2595. 10.1038/s41467-021-22822-9.

59. Chen, K., Henn, D., Januszyk, M., Barrera, J.A., Noishiki, C., Bonham, C.A., Griffin, M., Tevlin, R., Carlomagno, T., Shannon, T., et al. (2022). Disrupting mechanotransduction decreases fibrosis and contracture in split-thickness skin grafting. Sci Transl Med 14, eabj9152. 10.1126/scitranslmed.abj9152.

60. Yu, G.T., Ganier, C., Allison, D.B., Tchkonia, T., Khosla, S., Kirkland, J.L., Lynch, M.D., and Wyles, S.P. (2025). Mapping epidermal and dermal cellular senescence in human skin aging. Aging cell 24, e14358. 10.1111/acel.14358.

61. Zou, Z., Long, X., Zhao, Q., Zheng, Y., Song, M., Ma, S., Jing, Y., Wang, S., He, Y., Esteban, C.R., et al. (2021). A Single-Cell Transcriptomic Atlas of Human Skin Aging. Dev Cell 56, 383–397.e388. 10.1016/j.devcel.2020.11.002.

62. Liu, Z., Bian, X., Luo, L., Björklund Å, K., Li, L., Zhang, L., Chen, Y., Guo, L., Gao, J., Cao, C., et al. (2024). Spatiotemporal single-cell roadmap of human skin wound healing. Cell Stem Cell. 10.1016/j.stem.2024.11.013.

63. Hong, Z.X., Zhu, S.T., Li, H., Luo, J.Z., Yang, Y., An, Y., Wang, X., and Wang, K. (2023). Bioengineered skin organoids: from development to applications. Mil Med Res 10, 40. 10.1186/s40779-023-00475-7.

64. Han, Y., Micklem, G., and Kim, S.Y. (2023). Transcriptional landscape of oncogene-induced senescence: a machine learning-based meta-analytic approach. Ageing Res Rev 85, 101849. 10.1016/j.arr.2023.101849.

65. Chen, X., Lu, Y., Wang, L., Ma, X., Pu, J., Lin, L., Deng, Q., Li, Y., Wang, W., Jin, Y., et al. (2023). A fast chemical reprogramming system promotes cell identity transition through a diapause-like state. Nat Cell Biol 25, 1146–1156. 10.1038/s41556-023-01193-x.

66. Mahmoudi, S., Mancini, E., Xu, L., Moore, A., Jahanbani, F., Hebestreit, K., Srinivasan, R., Li, X., Devarajan, K., Prélot, L., et al. (2019). Heterogeneity in old fibroblasts is linked to variability in reprogramming and wound healing. Nature 574, 553–558. 10.1038/s41586-019-1658-5.

67. Milanovic, M., Fan, D.N.Y., Belenki, D., Däbritz, J.H.M., Zhao, Z., Yu, Y., Dörr, J.R., Dimitrova, L., Lenze, D., Monteiro Barbosa, I.A., et al. (2018). Senescence-associated reprogramming promotes cancer stemness. Nature 553, 96–100. 10.1038/nature25167.

68. Ruifrok, A.C., Katz, R.L., and Johnston, D.A. (2003). Comparison of quantification of histochemical staining by hue-saturation-intensity (HSI) transformation and color-deconvolution. Applied immunohistochemistry & molecular morphology : AIMM 11, 85–91. 10.1097/00129039-200303000-00014.

