## Supplementary figures and images for "Senescence-induced reparative fibroblasts enable scarless wound healing in aged murine skin"

### supplemental figures

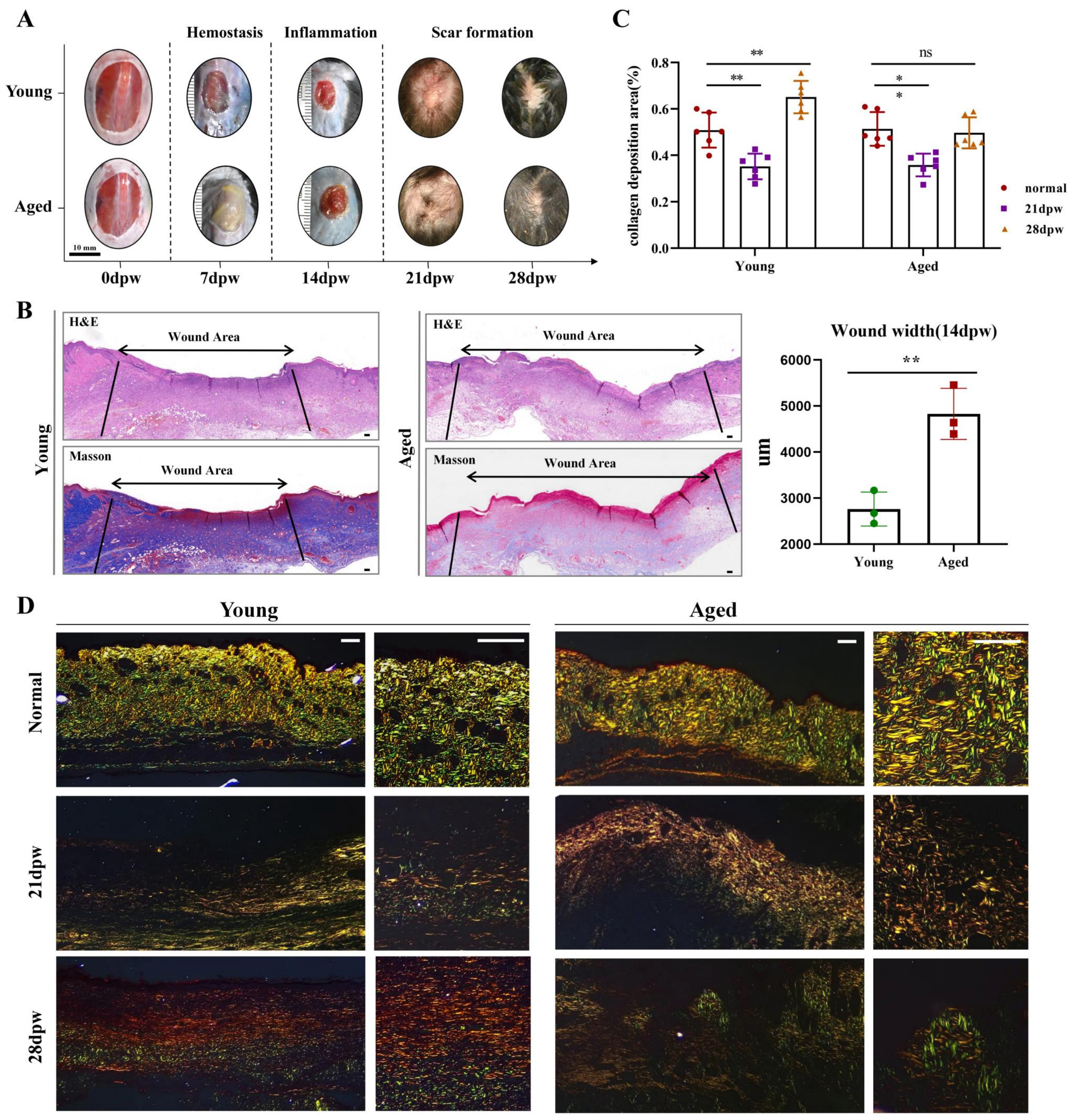

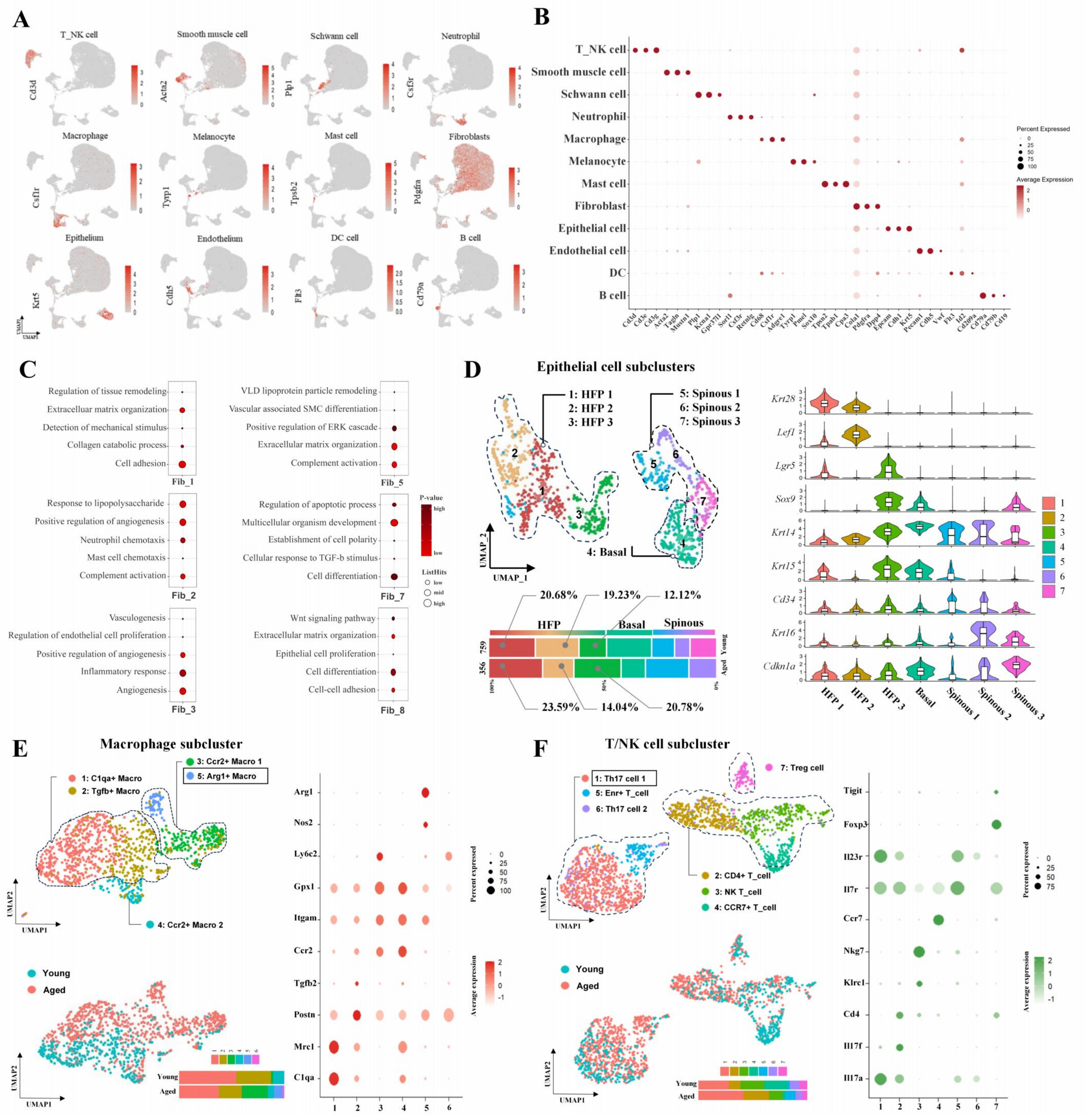

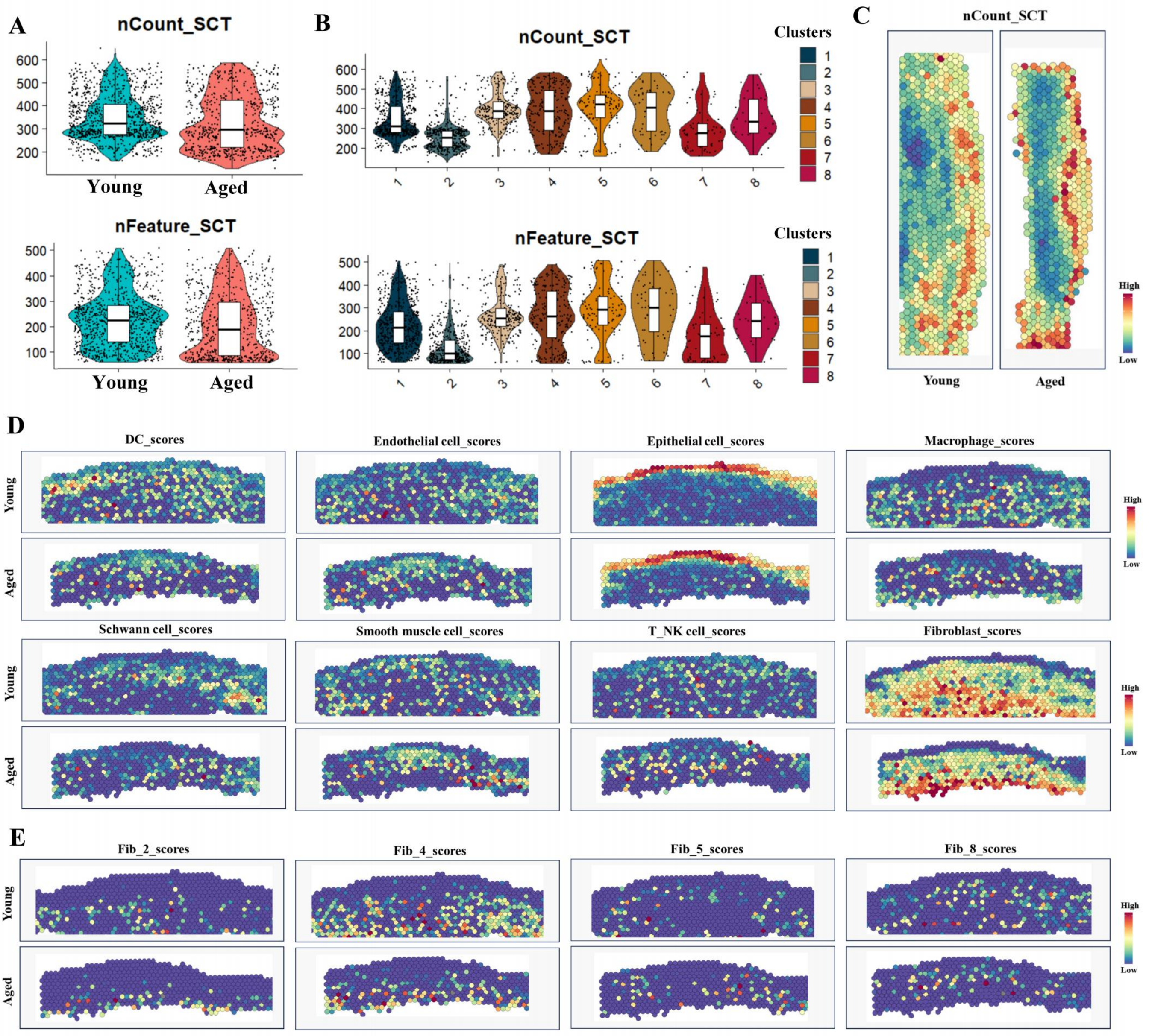

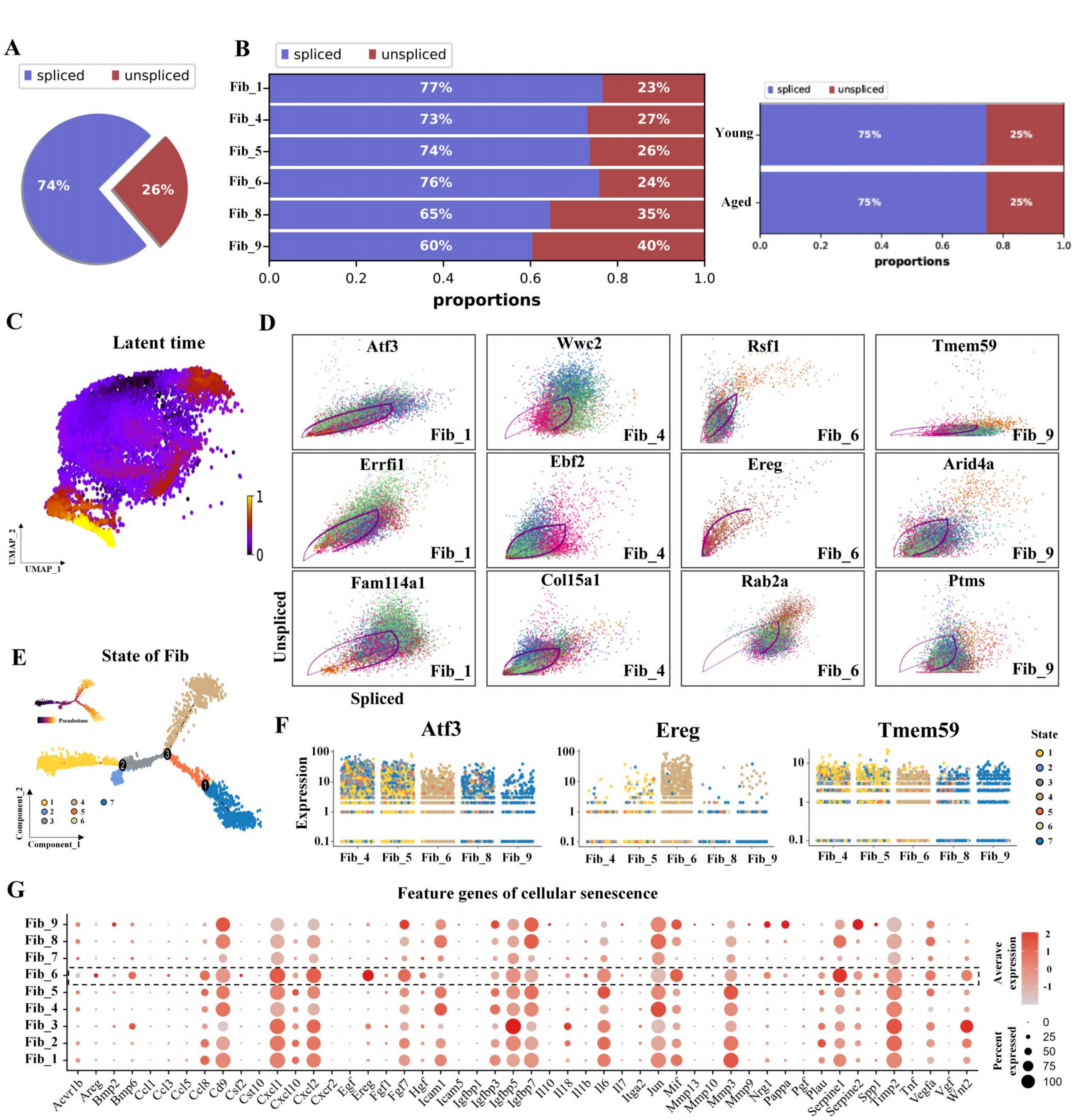

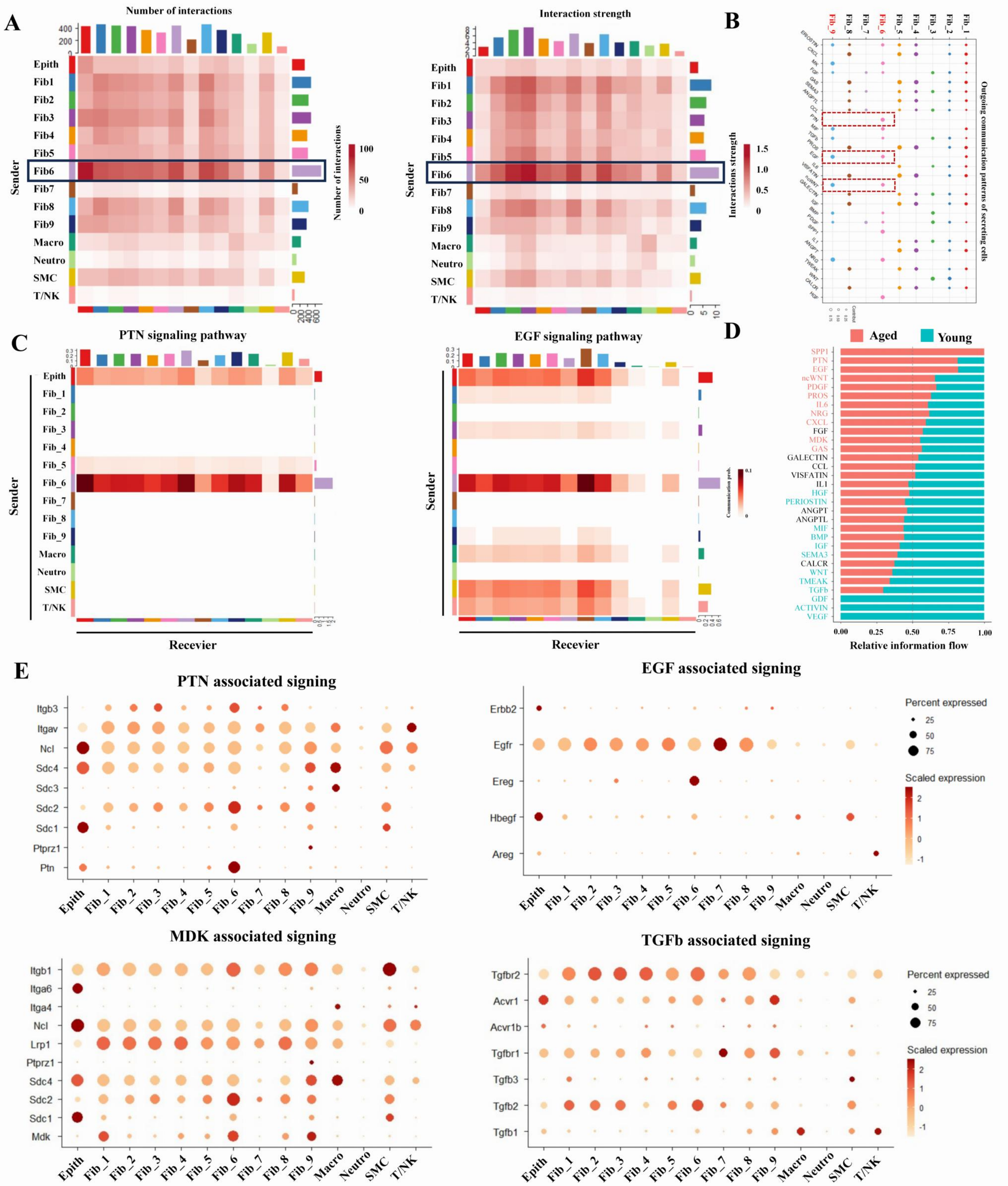

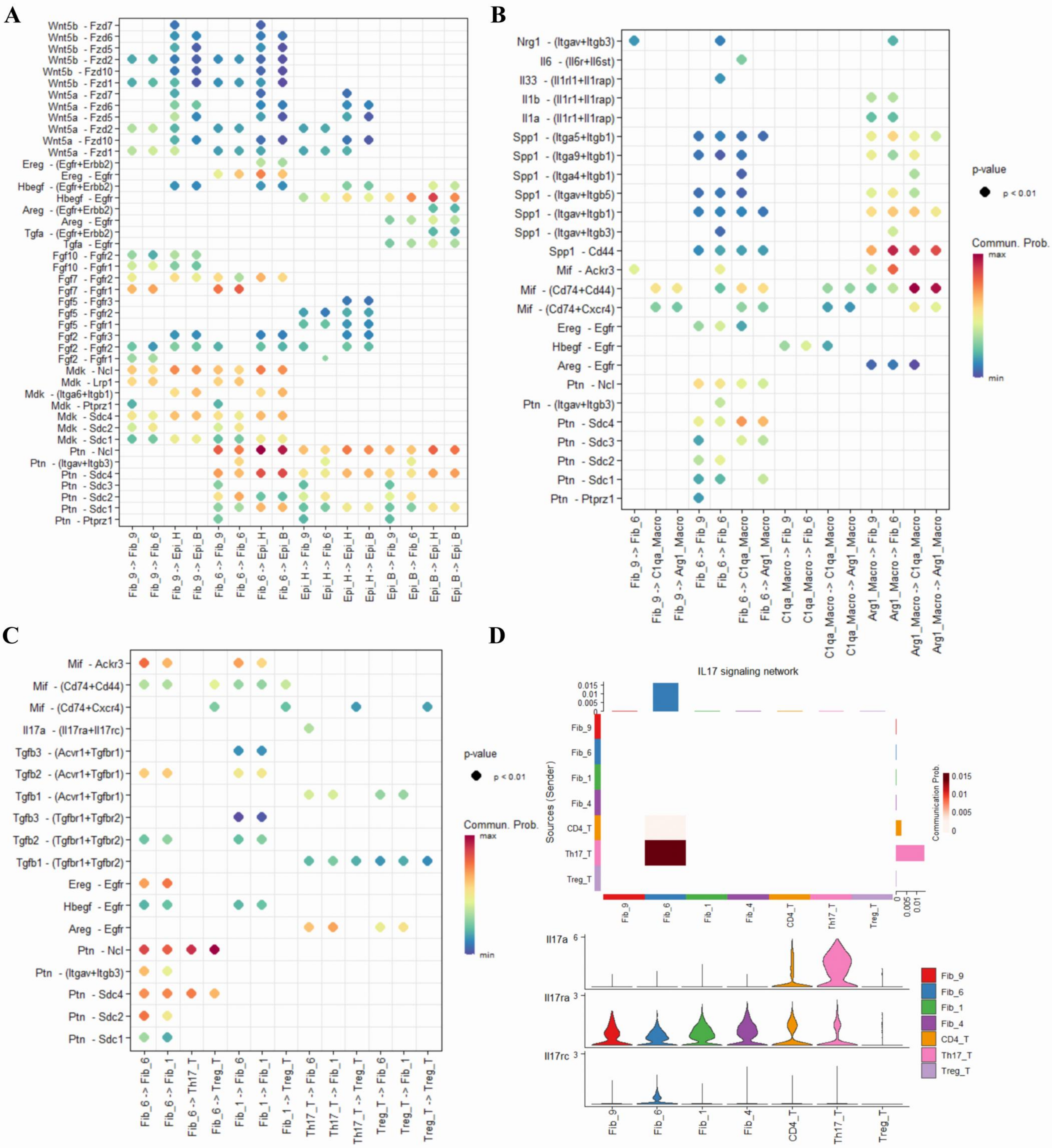

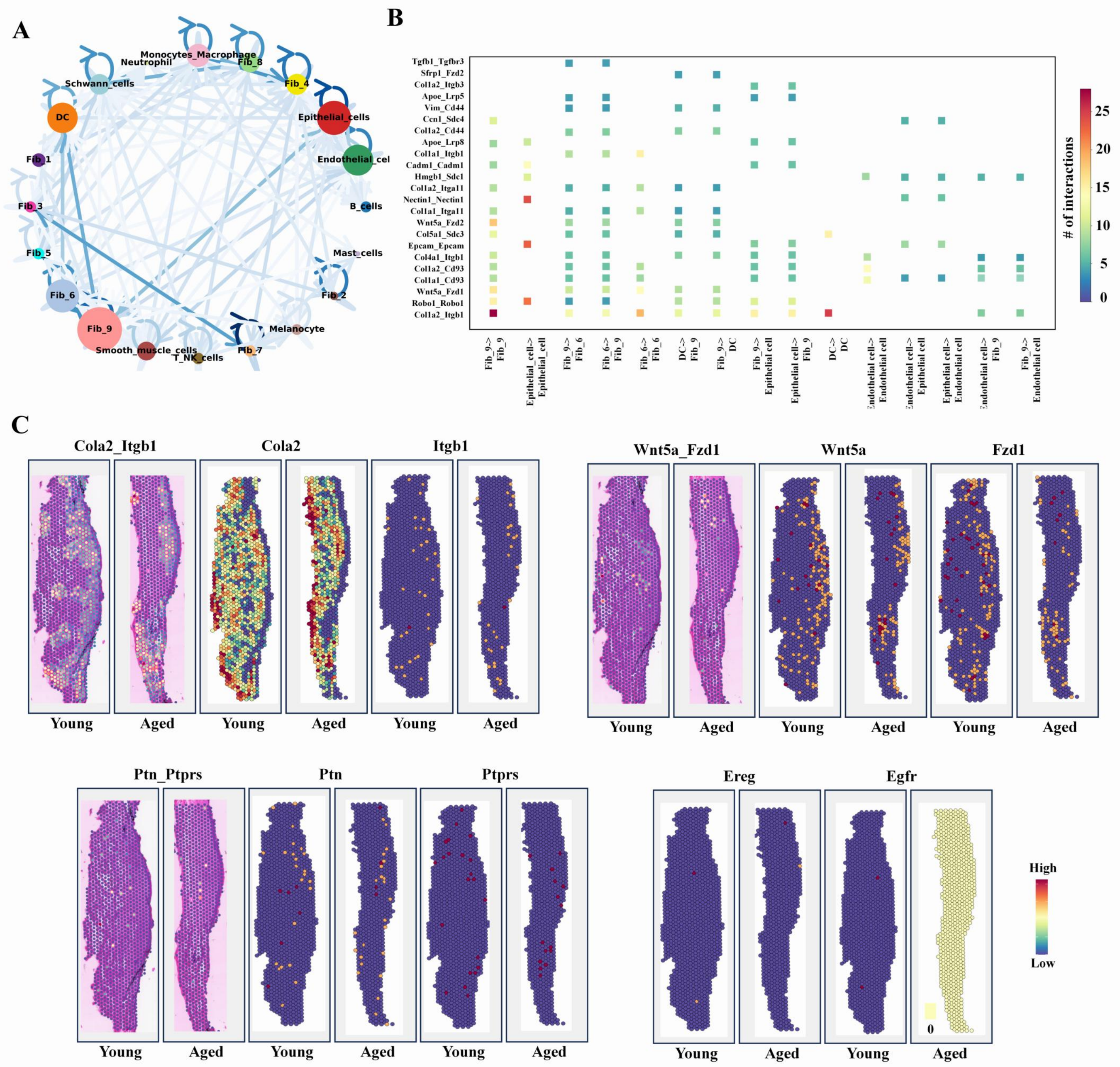

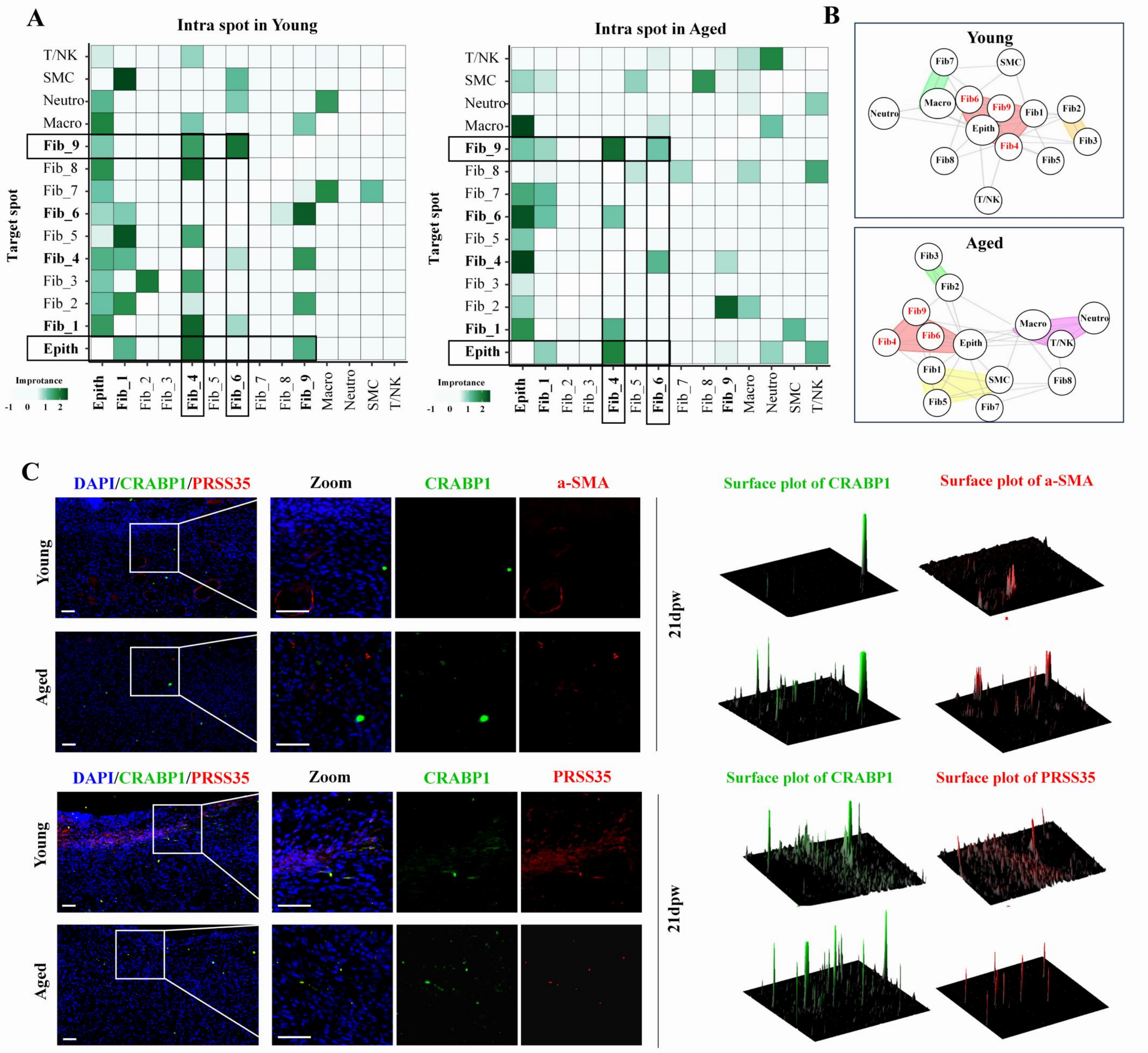

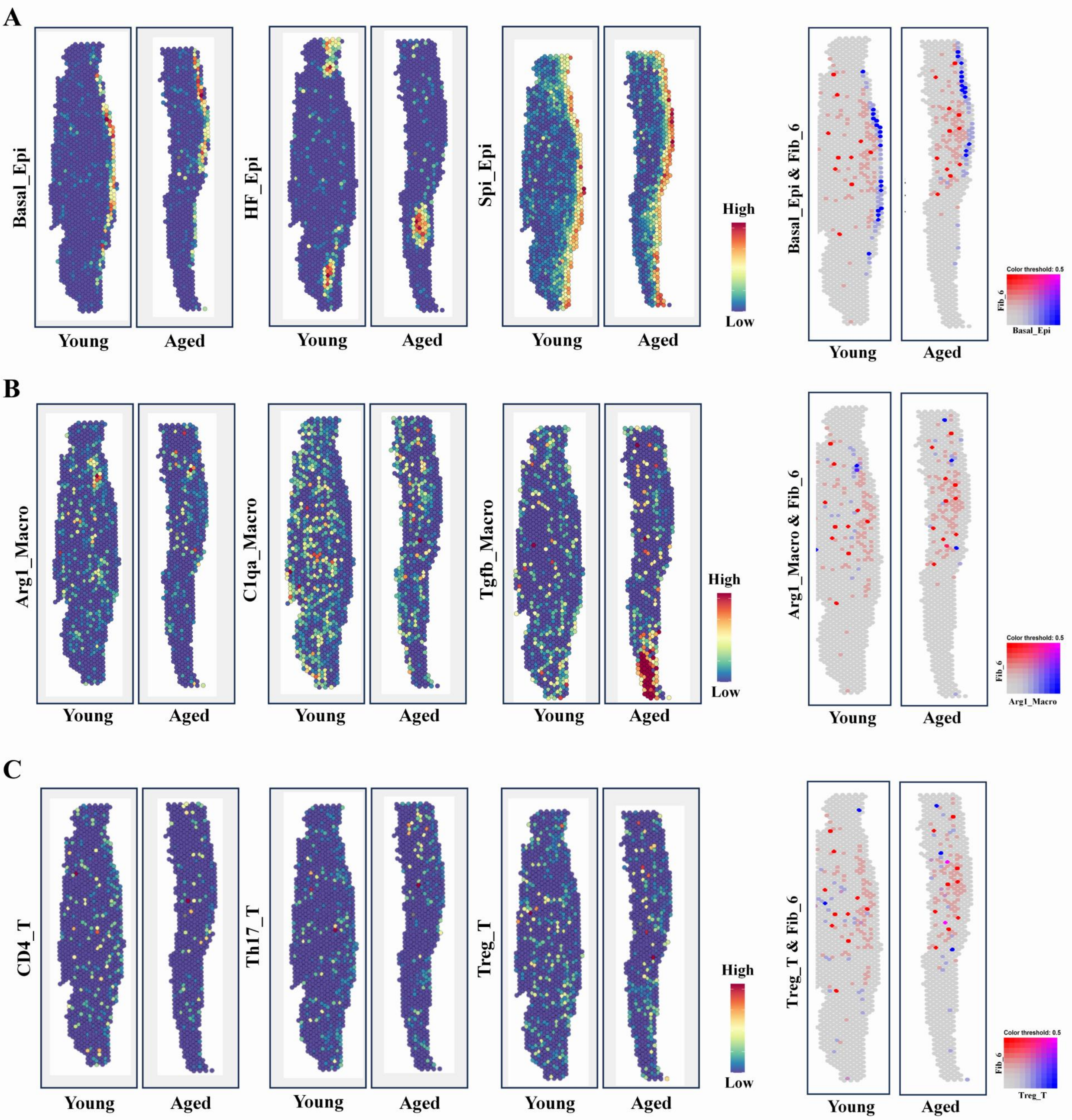

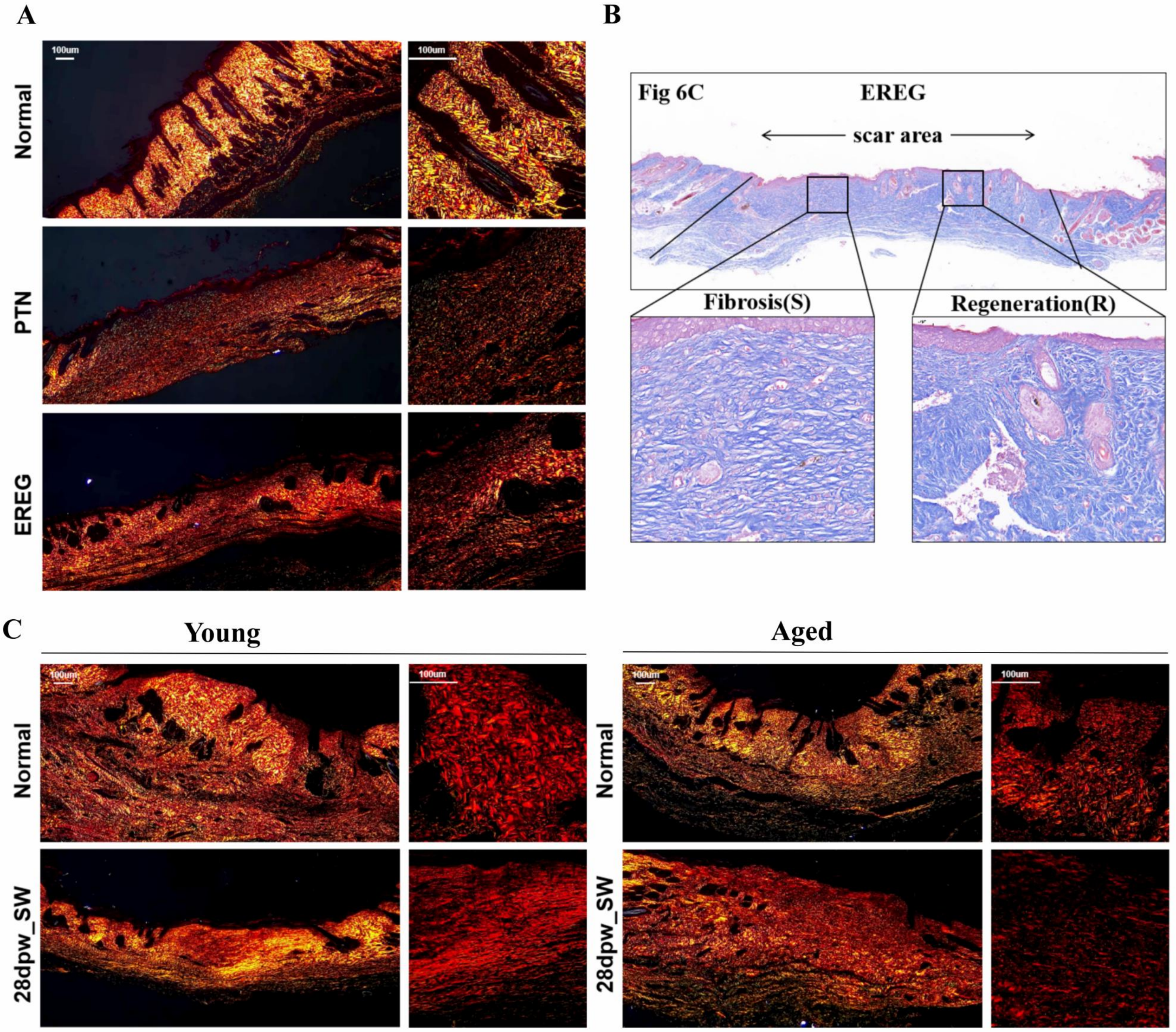
