## Supplementary material for "Senescence-induced reparative fibroblasts enable scarless wound healing in aged murine skin": Legends of supplemental figures

Supplementary Fig 1:

(A) Representative photos of the wound healing process at four phases. Scale bar as listed in the image.

(B) Hematoxylin-eosin (H&E) and Masson staining of wound area at 14 days post-wounding (dpw) in young and aged groups (left). Scale bar: 100 μm. Wound width differences were analyzed by bidirectional t-tests (n=3, **p < 0.01) (right).

(C) Quantification analysis for the ratio of collagen deposition area in scar tissues and normal skin assessed by bidirectional t-tests (n=5, **p < 0.01, ns, no significance).

(D) Representative Sirius red staining images of scar tissues and normal skin. Scale bar, 100um.

Supplementary Fig 2:

(A) UMAP plots showing the distribution of twelve cell types in single-cell RNA sequencing (scRNA-seq).

(B) Dot plot displaying canonical gene markers for each cell type. Dot size represents the percentage of cells expressing the gene; color indicates the average expression level.

(C) Dot plot showing gene ontology enrichment terms of fibroblast subclusters.

(D) Subclustering of Epithelial cells showing proportions(lower left) and gene marker (right) of seven subclusters. HFP, hair follicle progenitors.

(E-F) Subclustering of Macrophage(E) and T/NK cells. UMAP plots showing proportion and distribution within groups and subclusters. Dot plots showing gene markers of respective subcluster. Solid line squares represented aged-specific subclusters.

Supplementary Fig 3:

(A-C) Vlnplots showing the number of gene counts based on the SCTransform algorithm within groups(A) and subclusters(B). Spatial feature plot showing the distribution of the number of gene counts(C).

(D) Spatial feature plot showing the distribution of major cell types based on cell deconvolution result assessed by RCTD algorithm.

(E) Spatial feature plot showing the distribution of characteristics fibroblast subclusters.

Supplementary Fig 4:

(A-B) Pie plot(A) and bar plots(B) showing the ratio of spliced/unspliced RNA in fibroblasts within groups and subclusters.

(C) UMAP plot showing differentiation latent time of characteristics fibroblast subclusters. Red represented the end differentiation state while purple represent the opposite.

(D) Differentiation driver genes of characteristics fibroblast subclusters based on RNA velocity analysis. Clusters above the dotted line represented the gene inductive state while the below represented the suppressive state. Colors identified the fibroblast subclusters.

(E-F) Monocle2 pesudotime analysis of the differentiation state(E) and change trend of key gene expression (F) in characteristics fibroblasts subclusters.

The expression level of cellular senescence-associated genes in fibroblast subclusters.

Supplementary Fig 5:

(A) Heatmap plots showing the number (left) and strength (right) of interactions in cell-cell communication. Color block represented sender cells(Y-axis) and receiver cells(X-axis).
(B) Dotplot showing outgoing communication patterns of Fibroblasts subclusters. The size of the dots represented cellular contribution to the signaling pathway. The dotted box signed the common communication patterns of Fib_6 and Fib_9.
(C) Heatmaps showing interaction strength of pleiotrophin (PTN, left) and epidermal growth factor (EGF, right) signaling pathways in cell-cell communication. Color block represented sender and receiver cells the same as (A).
(D) Relative information flow showing up-regulated signaling between young and aged groups. Red, Aged specific. Blue, Young specific. The black signaling represents no differences.
(E) Dot plots showing the expression level of corresponding genes in PTN/EGF/MDK/TGFb signaling.

Abbreviation: Epith, epithelial cell. Macro, macrophage. SMC, smooth muscle cell. Neutro, neutrophil. T/NK, T/NK cells.

Supplementary Fig 6:

(A) Bubble plot showing ligand and receptor pairs between fibroblasts and epithelial subclusters.
(B) Bubble plot showing ligand and receptor pairs between fibroblasts and macrophage subclusters.
(C) Bubble plot showing ligand and receptor pairs between fibroblasts and T/NK cell subclusters.
(D) Heatmap plots showing interaction strength of IL17 signaling in communication of fibroblasts and T/NK cell subclusters (upper). Vlnplot showing expression level of corresponding genes in IL17 signaling (lower).

Abbreviation: Epi_B/Epi_H, basal/hair follicle progenitors epithelial cells. Macro, macrophage. CD4_T/Treg/Th17_T, CD4+/Treg/Th17 T cells.

Supplementary Fig 7:

(A) Number of cell-cell communication in spatial transcriptomics using CCI analysis.
(B) Dot plot showing ligand and receptor interactions assessed by CCI analysis. The x-axis represented sender->receiver cells.
(C) Spatial feature plot showing the expression level of ligand and receptor pairs.

Supplementary Fig 8:

(A) Heatmap plots showing the associations within the intra spot in ST using the MistyR algorithm, with the color of importance interpreted as spatial dependency.
(B) The communities plots showing spatial interactions (gray line) and proximity (polygon) within the intra-spot.
(C) Immunofluorescence staining for CRABP1 (green, marker of papilla fibroblast), a-SMA (red, marker of myofibroblast), and PRSS35 (red, marker of reparative fibroblast) in the scar tissues at 21 dpw(left). Surface plot showing relative fluorescence intensity of marker protein(right). Scale bar, 50 um. Dpw, day post wound.

Abbreviation: SMC, smooth muscle cell. Epith, epithelial cell. Neutro, neutrophil. T/NK, T/NK cells. Macro, macrophage.

Supplementary Fig 9:

(A) Spatial distribution of basal epithelial cells (Basal_Epi), hair follicle epithelial cells (HF_Epi) and spinous cells (Spi_Epi) and colocalization between Basal_Epi and fibroblast subcluster (Fib_6)
(B) Spatial distribution of macrophage subclusters(Arg1+, C1qa+, Tgfb+) and colocalization between Arg1+ macrophage and fibroblast subcluster (Fib_6).
(C) Spatial distribution of T/NK cell subclusters (CD4+, Th17, Treg) and colocalization between Treg and fibroblast subcluster (Fib_6).

Supplementary Fig 10:

(A) Representative Sirius red staining images of normal skin and scar tissues in PTN and EREG treatment group. Scale bar, 100um.

(B) Hematoxylin-eosin (H&E) and Masson staining for regeneration and fibrosis area in EREG treatment samples. Scale bar, 200um.

(C) Representative Sirius red staining images of scar tissues in small wounds (SW) and normal skin. Scale bar, 100um. Dpw, day post wound.
